# Hypoxic stress dysregulates functions of glioma-associated myeloid cells through epigenomic and transcriptional programs

**DOI:** 10.1101/2024.09.12.612769

**Authors:** Monika Dzwigonska, Patrycja Rosa, Beata Kaza, Szymon Lipiec, Salwador Cyranowski, Aleksandra Ellert-Miklaszewska, Agata Kominek, Tomasz Obrebski, Anna R. Malik, Katarzyna Piwocka, Jakub Mieczkowski, Bozena Kaminska, Katarzyna B. Leszczynska

**Affiliations:** Laboratory of Molecular Neurobiology, Nencki Institute of Experimental Biology Polish Academy of Sciences, Pasteura St. 3, 02-093 Warsaw, Poland; Laboratory of Cytometry, Nencki Institute of Experimental Biology Polish Academy of Sciences, Pasteura St. 3, 02-093 Warsaw, Poland; Cellular Neurobiology Research Group, Institute of Developmental Biology and Biomedical Sciences, Faculty of Biology, University of Warsaw, Miecznikowa St. 1, Warsaw 02-096, Poland; 3P-Medicine Laboratory, Medical University of Gdansk, 80-210 Gdansk, Poland

**Keywords:** tumour-associated myeloid cells, glioma, microglia, macrophages, hypoxia, epigenetics, chromatin accessibility

## Abstract

Hypoxia rapidly alters gene expression to allow cellular adaptation to challenging conditions and support tumour growth. Hypoxia also affects the chromatin structure by modifications of histones and DNA methylation. Glioblastoma (GBM) is an aggressive, deadly primary brain tumour for which there is no effective treatment. The tumour microenvironment of GBM is highly heterogeneous, with infiltration of glioma-associated microglia and macrophages (GAMs) and the presence of necrotic, hypoxic regions which significantly impair effectiveness of therapies. The mechanisms through which hypoxia alters the tumour microenvironment and regulates functions of infiltrating immune cells remain poorly understood.

Here, we show that hypoxia modulates the expression of myeloid markers in distinct ways: upregulates the expression of monocytic marker *Lgals3* and downregulates the microglial markers *P2ry12* and *Tmem119* in microglial and monocytic GAMs *in vitro* and *in vivo*. Underlying genome-wide hypoxia-dependent transcriptomic changes in microglial cells were identified using microglia-glioma co-cultures and validated in human and mouse GBM single-cell transcriptomics datasets. Numerous GAM subtype markers are dysregulated in response to hypoxic stress due to associated changes in chromatin accessibility, as determined using ATACseq. While hypoxia alone drives a decrease of the overall chromatin accessibility at gene promoters, the exposure to glioma cells under hypoxic conditions leads to both increases and decreases of chromatin accessibility at promoter regions in microglial cells. Hypoxia downregulates the chromatin accessibility at the regions enriched in motifs for transcription factors known as master regulators of microglial cell identity and function, including *SPI1* or *IRF8*. Overall, our results highlight the importance of hypoxic stress as a strong intratumoral regulator of myeloid cell functions, which adds a new dimension to the characterisation of particular GAM subpopulations.

## INTRODUCTION

Glioblastoma (GBM) is the most aggressive primary brain tumour, incurable with existing therapies ^1^. The highly heterogeneous tumour microenvironment (TME) significantly contributes to the complexity and malignancy of GBM ^2^. Myeloid cells recruited to the tumour, including brain-resident microglia (Mg) and bone marrow-derived monocytes and macrophages (BMDMs), collectively known as tumour- or glioma-associated microglia/macrophages/monocytes (TAMs or GAMs, respectively), constitute up to 30% of the GBM mass ^3,4^. Recent applications of single-cell transcriptomics characterised GAM subpopulations in human GBMs and experimental gliomas, revealing a heterogeneity of myeloid cells ^5–12^. The two major groups: microglia-GAMs (Mg-GAMs) and monocytic/macrophage GAMs (Mo/Mφ-GAMs) were further stratified into specialised subpopulations with predominant phenotypes: homeostatic, interferon, chemotactic, phagocytic, lipid, ribosomal, hypoxic, proliferative or transitory phenotypes ^5–8,12,13^. These subpopulations of GAMs have specific gene expression patterns, which are often used to select the characteristic markers for each GAM subtype. Despite some functional specialisation, most GAM subpopulations actively support tumour progression and immunosuppression ^3,14^.

Advances in spatial transcriptomics showed that the proportions of Mg-GAMs and Mo/Mφ- GAMs are different in certain tumour areas, with an indication that a higher proportion of Mo/Mφ-GAMs is found at the tumour core, potentially due to an influx of monocytes through leaky, disorganised tumour vasculature at the later tumour stages ^7,8,10,15^. Mg-GAMs, on the other hand, are more enriched at the tumour periphery, tumour border and the surrounding brain parenchyma ^5,6,14^. While in the tumour core, GAMs can be exposed to hypoxic stress (shortage of oxygen), which leads to rapid gene expression changes to allow adaptation to challenging oxygen-deprived conditions ^16,17^. However, the role of hypoxia in reprogramming of GAMs is only starting to emerge. Some studies showed that while in hypoxia, GAMs acquire immunosuppressive and immunotolerant phenotypes ^14^ or can increase the permeability of GBM vasculature through the release of adrenomedullin ^8^. Pre-clinical testing demonstrated that targeting adrenomedullin in GBM is a good strategy to improve blood vessel normalisation and drug delivery ^8^. A recently discovered new subtype of GAMs, “lipid-laden TAMs”, were found to reside in hypoxic niches and use myelin-derived lipids to fuel the aggressiveness of glioma cells ^11^. Altogether, these findings emphasize the importance of hypoxic stress in shaping distinct GAM phenotypes in GBM. However, it remains unclear if and how hypoxia can shift GAM subpopulations into particular states and phenotypes.

Single-cell transcriptomics data defined the sets of genes that are characteristic for particular Mo/Mφ-GAM and Mg-GAM sub-clusters ^5–8,12^. However, some myeloid markers, which are highly expressed across multiple subtypes of GAMs, are affected by various environmental cues, which raises the awareness about usefulness of selective GAM markers to detect a particular GAM subpopulation, e.g. in immunohistochemistry. For example, *LGALS3,* the gene encoding galectin-3 (GAL3) is often used as a general monocytic and macrophage GAM marker distinguishing the Mo/Mφ-GAMs from Mg-GAMs ^5,8^. However, *LGALS3* is also highly expressed in more specialised lipid Mo/Mφ-GAMs due to its function in lipid processing and accumulation ^6,8^. In addition, some studies showed that *LGALS3* is hypoxia-inducible in a number of cell types, including cancer cells ^18,19^. Therefore, we hypothesised that *LGALS3,* and potentially some other myeloid subtype markers, could be altered by hypoxic stress either in Mg-GAMs or in Mo/Mφ-GAMs, which in turn could affect the assessment of the spatial distribution of various GAM subpopulations. The mechanisms underpinning transcriptomic changes driven by hypoxic stress in GAMs are poorly defined. Here, we showed that along with known hypoxia-regulated genes *Vegfa* and *Glut1* ^20,21^, hypoxia upregulates the monocytic/macrophage marker *Lgals3* expression and downregulates the microglial markers *P2ry12* or *Tmem119*, both in microglial and macrophage cells. We find similar correlations between the hypoxic score and gene expression *in vivo* in human and mouse GBM single-cell transcriptomics datasets. Using direct microglia-glioma co-culture we defined genome-wide hypoxia-dependent transcriptomic changes in microglial cells. We found that numerous GAM subtype markers were dysregulated in response to hypoxic stress, and these changes were associated with epigenomic alterations detected with ATACseq. While hypoxia alone decreases the overall chromatin accessibility at gene promoters, contacts with glioma cells under hypoxic conditions lead to far more extensive remodelling of chromatin accessibility at promoter regions in microglial cells. Our data highlights hypoxic stress as a strong intratumoral regulator of myeloid subtype gene expression by imposing epigenomic and transcriptomic changes.

## RESULTS

### Hypoxia dysregulates expression of *Lgals3*, *P2ry12* and *Tmem119* in myeloid cells *in vitro* and in GAMs *in vivo*

We investigated whether hypoxia directly alters *Lgals3* expression in various macrophage and microglial cells using mouse microglial BV2 cells, RAW 264.7 cells (a tumour-derived mouse macrophage cell line) and primary mouse bone marrow-derived macrophages (BMDM). In three cell types, hypoxic conditions (<0.1% O2) led to upregulation of *Lgals3* expression at the mRNA and protein levels, along with known hypoxia-induced genes *Vegfa* and *Glut1 (Slc2a1)* (**Figure 1A-B**). *Lgals3* expression was also modulated in response to a milder hypoxia (1% O2) and a hypoxia-mimetic drug (CoCl2), which inhibits the activity of prolyl hydroxylases and therefore leads to HIF-α stabilisation in normoxic conditions ^22^. Both the mild hypoxia and CoCl2 upregulated the expression of *Lgals3* at the mRNA and protein levels (**Supplementary** Figure 1A-B). Interestingly, under the same conditions, the expression of microglial markers *Tmem119* and *P2ry12* was significantly downregulated. While *Tmem119* and *P2ry12* are microglia markers, their expression was detected in RAW 264.7 and BMDM cells, and was downregulated by hypoxia (**Figure 1C**). Moreover, the mild hypoxia and CoCl2 also decreased the expression of these genes (**Supplementary** Figure 1C). Overall, this data suggests that hypoxic stress regulates the expression of *Lgals3* and *Tmem119* and *P2ry12* in opposite directions, irrespective of the origin of the myeloid cells.

**Figure 1.**
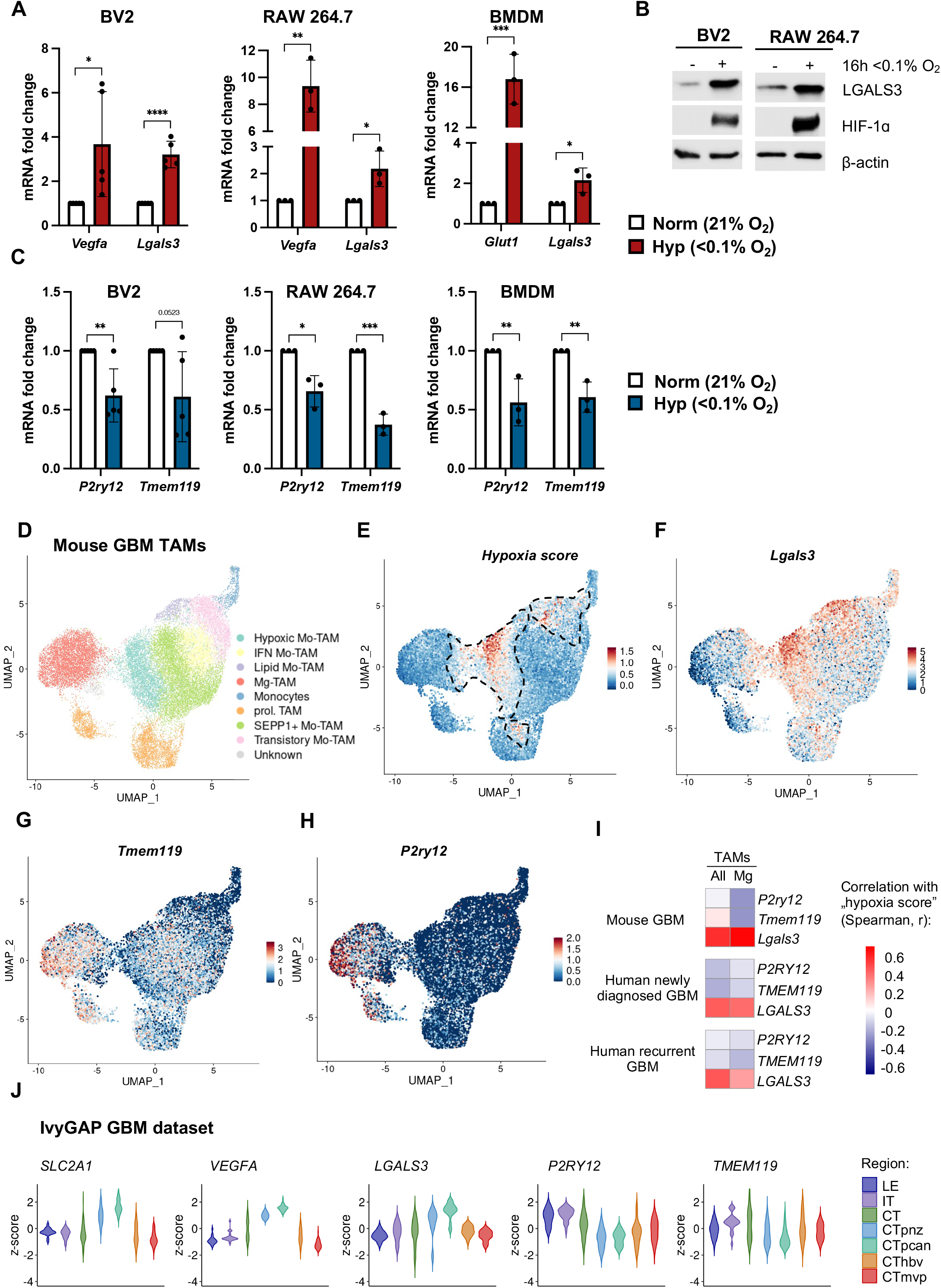
Hypoxia dysregulates the expression of *Lgals3, P2ry12* and *Tmem119* in myeloid cells *in vitro* and in GBM samples *in vivo*. **A,** Expression analysis using qPCR of *Lgals3* in BV2, RAW 264.6 and BMBM cells, as indicated in relation to *Rn18s* housekeeping gene. *Glut1* (*Slc2a1*) and *Vegfa* were used as hypoxia controls. The mean fold change with standard deviation (SD) is shown from at least three independent biological replicates. Two-tailed student t-tests test determined the statistical significance (* *p*<0.05, ** *p*<0.01, *** *p*<0.001). **B,** BV2 and RAW 264.7 cell lines were exposed to 21% or <0.1% O2 for 16 h and subjected to western blotting with the anti-LGALS3 antibody. HIF-1α was used as a hypoxia control. A representative experiment of three independent biological repeats is shown. **C,** Expression analysis using qPCR of *P2ry12 and Tmem119* in BV2, RAW 264.6 and BMDM cells, as indicated in relation to *Rn18s* housekeeping gene. The mean fold change with standard deviation (SD) and statistical analysis as in (**A**) is shown from at least three independent biological replicates. **D,** UMAP plot showing clusters of monocytes and TAM subpopulations from mouse GL261 tumours as identified in the public scRNA-seq dataset (GSE163120) ^6^.| **E,** UMAP plot showing the expression of “hypoxia score” in the dataset from (**D**). **F-H,** UMAP plots showing the expression of (**F**) *Lgals3*, (**G**) *Tmem119* and (**H**) *P2ry12* in the dataset from (**D**). **I,** Heatmaps showing the correlations between “hypoxia score” and *P2ry12*, *Tmem119* and *Lgals3* expression in mouse GL261 tumours, human newly diagnosed GBM and human recurrent GBM scRNA-seq Antunes-datasets (all data from GSE163120). **J,** Violin plots showing z-score expression of hypoxia-inducible genes *VEGFA* and *SLC2A1*, as well as *TMEM119*, *P2RY12 and LGALS3* genes in different anatomical regions of GBM samples from Ivy GBM Atlas Project dataset (Ivy GAP; https://glioblastoma.alleninstitute.org/) ^23^. LE, leading edge; IT, infiltrating tumour; CT, cellular tumour; CTpnz, perinecrotic zone; CTpcan, pseudopalisading cells around necrosis; CThbv, hyperplastic blood vessels in cellular tumour; CTmvp, microvascular proliferation.

We verified the expression of these genes in mouse GBM GAMs *in vivo* in relation to hypoxia, by exploring the public scRNAseq dataset from Antunes *at al.*, which has annotations of multiple GAM sub-clusters, including microglia, monocytes, lipid, proliferative, IFN, but also hypoxic clusters (in relation to this data we use nomenclature TAMs rather than GAMs, as in the Antunes publication) ^6^. The published annotation of clusters in this dataset shows that cells in the hypoxic cluster (**Figure 1D**) have high *Lgal3* expression, and that hypoxic cells are mostly assigned to the Mo-TAMs clusters rather than to the Mg-TAMs (**Figure 1D-F**). We extracted the top genes expressed in hypoxic TAMs cluster from another scRNAseq GBM dataset from Wang *et al.,* and created a combined “hypoxia score”, which encompasses the common top hypoxia TAM cluster genes from both studies (**Supplementary Table S1**) ^6,8^. We then analysed the expression of the combined “hypoxia score” in the scRNAseq Antunes- dataset. In addition to the initial hypoxic Mo-TAMs cluster (**Figure 1D**), some groups of hypoxic cells were also visible in other clusters of TAMs, particularly in the lipid Mo-TAMs, transitory Mo-TAMs, proliferative Mo-TAMs and in Mg-TAMs (**Figure 1E**). In agreement with the literature, *Lgals3* expression was stronger in the whole Mo-TAM subpopulation in comparison to Mg-TAMs ^5^, however, it was also upregulated in TAMs with increased “hypoxia score” in proliferative, lipid, monocytic and transitory Mo-TAMs (**Figure 1F**). The expression of *Tmem119* and *P2ry12* was higher in Mg-TAMs than in Mo-TAMs, however, it decreased in Mg-TAMs along with increased “hypoxia score” (**Figure 1G-H**). Gene expression shows a statistically significant positive correlation of the “hypoxia score” with *Lgals3* and a negative correlation with *Tmem119* and *P2ry12* in TAMs, and particularly in Mg-TAMs (**Figure 1I**). Importantly, these observations were confirmed in the human newly-diagnosed and recurrent GBM TAM Antunes-datasets (**Figure 1I** and **Supplementary** Figure 2) ^6^.

To corroborate these findings further, we analysed the human GBM samples from the IvyGAP glioblastoma project, where the gene expression was tested in samples microdissected from the distinct spatial locations in GBM biopsies, including perinecrotic regions known to be hypoxic, vascularised areas, tumour leading edge, infiltrative tumour or cellular tumour ^23^. The expression of *LGALS3* was the highest in the perinecrotic and pseudopalisading regions (CTpnz and CTpcan) in a similar way as the expression of some typical hypoxic markers such as *SLC2A1* or *VEGFA* (**Figure 1J**). On the contrary, the expression of *TMEM119* and *P2RY12* was the lowest at the hypoxic perinecrotic and pseudopalisading areas and the most abundant in the infiltrating tumour (IT) and tumour leading edge (LE) (**Figure 1J**). While it may reflect the fact that *TMEM119* or *P2RY12* expressing cells are less abundant in the perinecrotic regions, this data also suggests that hypoxia strongly dysregulates the expression of the tested genes in tumours *in vivo*, irrespectively of the cell types exposed to hypoxic stress.

### Reprograming of the transcriptome of BV2 microglial cells by glioma and hypoxia

To better understand the role of hypoxia in controlling the gene expression in myeloid cells associated with the tumour, we have employed direct microglia-glioma co-cultures, which recapitulate tumour-microglia interactions during tumour progression ^24^. Glioma cells (GL261- GFP) were seeded in culture dishes and the BV2 microglial cells were added the next day (**Figure 2A**). After 24 hours, BV2-GL261 co-culture was either transferred into hypoxic conditions (<0.1% O2) or left in normoxia (21% O2) for additional 16 hours. BV2 cells seeded without glioma cells served as controls. After the hypoxic treatment, the co-cultures were dissociated into single-cell suspensions, fixed with glyoxal, and BV2 cells immuno-labelled with anti-CD45-PE antibody were separated from glioma cells by FACS sorting, which allowed obtaining pure populations of cells (**Supplementary** Figure 3A-B). The glyoxal fixative was used straight after the hypoxic treatment to prevent any reoxygenation-dependent gene expression changes that could occur during cell staining and FACS procedure steps. The glyoxal fixation was recently validated as suitable for the downstream RNAseq analysis ^25^, and we also confirmed the high RNA quality of glyoxal-fixed samples, and induction of hypoxia-dependent genes by qPCR (**Supplementary** Figure 3C-D). Moreover, we observed morphological changes in BV2 microglial cells co-cultured with glioma cells, characterised by more elongated shapes in hypoxic conditions (**Figure 2B**).

**Figure 2.**
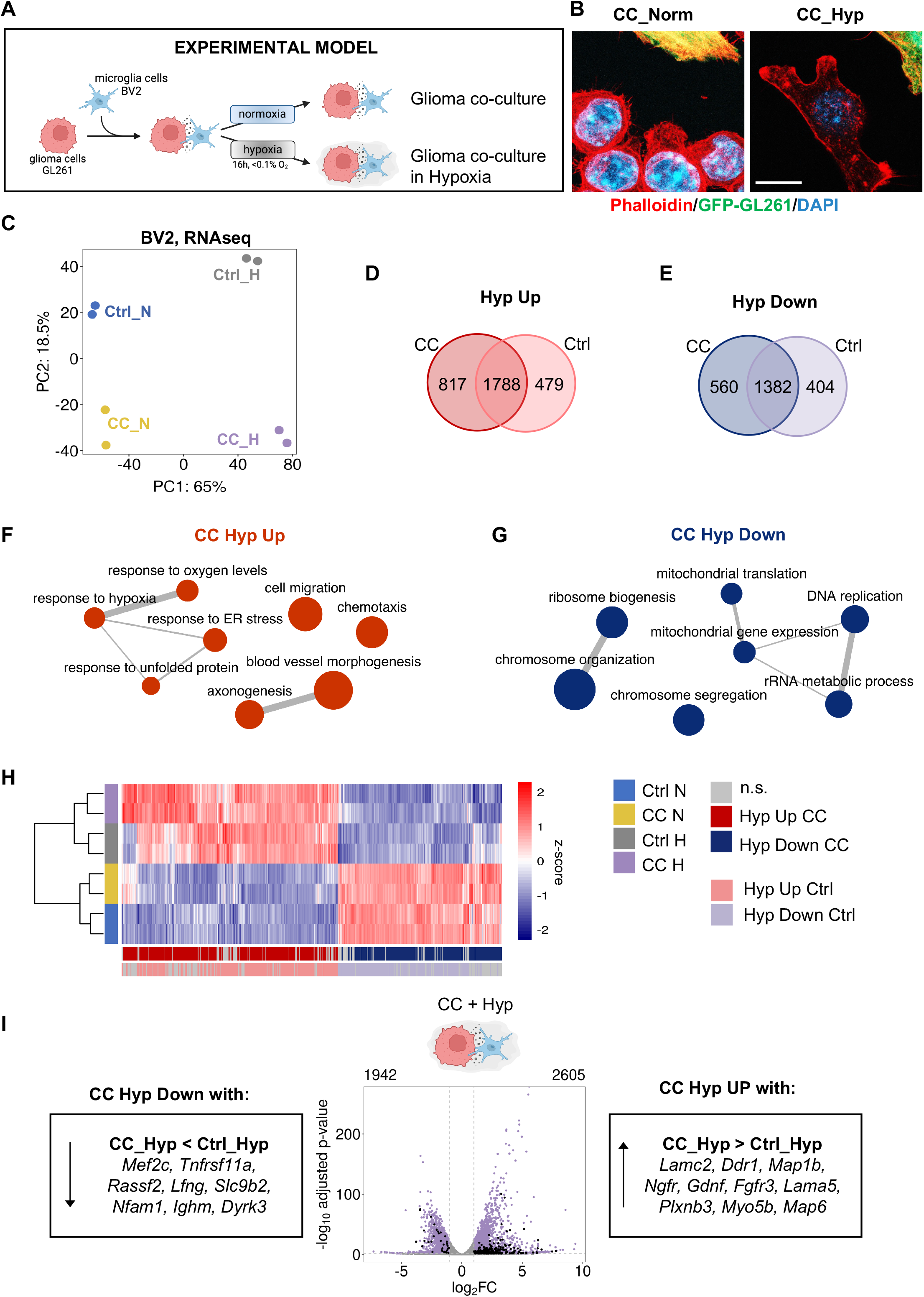
Glioma and hypoxia exposure reprograms the transcriptome of BV2 microglial cells. **A,** A scheme summarising direct glioma-microglia co-culture experiment. **B,** Immunofluorescent images of BV2 and GL261(stably expressing GFP) cells co-cultured in normoxic or hypoxic conditions as in (**A**). Red colour visualises actin cytoskeleton stained with phalloidin and blue colour marks nuclei stained with DAPI. Green to yellow cells represent GL261 while red-only positive cells are microglia cells, which are negative for GFP. Scale bar, 10 μm. **C,** A principal component plot of normalised gene expression in BV2 cells in normoxia (Ctrl_N), in hypoxia (Ctrl_H) or in normoxic co-cultures with glioma (CC_N) and in hypoxic co-cultures with glioma (CC_H). **D-E,** Venn diagrams summarising the overlap between differentially upregulated (**D**) and downregulated (**E**) genes in BV2 cells after exposure to hypoxia in the presence (CC) or absence (Ctrl) of glioma cells. **F-G,** Dot plots of enriched GO pathways for gene sets significantly upregulated (**F**) or downregulated (**G**) after exposure to hypoxia in coculture with glioma cells (CC). The top 15 GO pathways were selected based on the lowest adjusted *p*-values, and closely related terms were merged using REVIGO. **H,** A heatmap showing z-score expression for genes up- or downregulated in CC or Ctrl hypoxic cells. Genes which were additionally increased or decreased in CC versus Ctrl hypoxic conditions are projected in light red and light blue, respectively. **I,** A volcano plot showing significantly up- (2606) or downregulated (1942) genes in glioma co-cultured (CC) microglial cells in purple. Genes with expression additionally increased or decreased in CC versus Ctrl hypoxic conditions are projected in black. Left and right box panels show examples of genes additionally decreased or increased in CC versus Ctrl, respectively.

The principal component (PC) analysis of the BV2 RNAseq data showed a clear separation between all four conditions tested: BV2 cells isolated from normoxic co-cultures with glioma (CC_N), BV2 cells isolated from hypoxic co-cultures with glioma (CC_H), BV2 monocultures in normoxia (Ctrl_N) and BV2 monocultures in hypoxia (Ctrl_H) (**Figure 2C**). A stronger separation of samples was observed due to hypoxic stress (samples separating along the PC1 axis) as opposed to the co-culture effect (samples separating along the PC2 axis). Differential gene expression analysis showed that co-culture of BV2 cells with glioma cells induced expression of 412 genes and decreased 44 genes (CC_N versus Ctrl_N; threshold at log2FC = 1 and p-value < 0.05; **Supplementary** Figure 4A). Exposure of BV2-GL261 co-cultures to hypoxia induced many more transcriptomic changes in BV2 cells in comparison to the normoxic co-culture, with 2605 upregulated and 1942 downregulated genes (CC_H versus CC_N; **Supplementary** Figure 4B). Hypoxia alone also induced a large number of transcriptomic changes in BV2 cells (2267 up and 1786 down; Ctrl_H versus Ctrl_N) (**Supplementary** Figure 4C), of which the majority overlapped with the gene expression changes induced by hypoxia in co-cultures (**Figure 2D-E**). Interestingly, genes that were up- or downregulated by hypoxia (either in co-cultures or alone) had a much higher mean basal expression than the genes that were altered in response to the contacts with glioma alone in normoxic conditions (**Supplementary** Figure 4D-F).

We confirmed that BV2 cells in monoculture or in co-culture with glioma cells showed upregulation of HIF activity upon exposure to hypoxia by measuring the expression of genes from the validated specific HIF signature (HIF metagene) ^26^. The HIF signature gene expression was significantly increased under hypoxic conditions (**Supplementary** Figure 4G). Moreover, the analysis of promoters of the upregulated genes showed the enrichment in the motifs for transcription factors (TFs) that are typical for the hypoxic response, including HIF-1α, HIF-2α, KLFs, ATF3 or ATF4 (**Supplementary** Figure 4H) ^26–30^. Overrepresentation analysis (ORA) of Gene Ontology (GO) terms using genes upregulated in glioma-co-cultured BV2 cells exposed to hypoxia showed the enrichment of the processes typical for hypoxia, including response to decreased oxygen levels, angiogenesis, blood vessels morphogenesis, chemotaxis or cell motility-related processes (**Figure 2F** and **Supplementary Table S2**). Interestingly, the upregulation of pathways related to axon development, axon guidance or tissue morphogenesis in BV2 microglial cells indicated a strong shift towards the changes in cell morphology (**Figure 2F**), which was in line with our observations from immunofluorescent analysis of the BV2 cell morphology in hypoxic co-culture (**Figure 2B**). The downregulated genes in BV2 cells from glioma co-cultures under hypoxia were associated with the pathways previously reported as repressed in hypoxia, including ribosomal biogenesis, RNA processing or DNA replication (**Figure 2G**) ^31,32^.

Further analysis of genes induced more significantly in the presence of hypoxia and glioma co-culture as opposed to the hypoxia alone, indicated processes related to axonogenesis, axon development and other morphological changes (**Figure 2H** and **Figure 2I**, right panel includes examples of genes from the axon development pathway; see **Supplementary Table S2** for the full list). Interestingly, the genes more decreased by hypoxia and co-culture with glioma cells in comparison to hypoxia alone included genes from the pathways related to leukocyte or myeloid cell differentiation, including *Mef2c, Tnfrsf11a, Rassf2, Lfng, Slc9b2, Nfam1, Ighm* or *Dyrk3* (**Figure 2I**, left panel) and **Supplementary Table S2**). This finding supports the hypothesis that hypoxia potentially influences the expression of myeloid marker genes in GAMs, *in vivo* hypoxia additionally imposes the gene expression changes driven by interactions of GAMs with glioma cells.

### Hypoxia dysregulates the expression of multiple Mo/Mφ and microglial marker genes

Based on the published scRNAseq data reports, which distinguished Mg-GAMs and Mo-GAMs populations, as well as functional subpopulations of Mo/Mφ -GAMs characterised as transitory, IFN, lipid, hypoxic, chemotactic, phagocytic, ribosomal or proliferative, we created a list of genes highly expressed in distinct GAM clusters across multiple studies (**Supplementary Table S3**) ^5–8^. We then investigated the expression of these genes in hypoxic BV2 cell monocultures (Hyp) or BV2 co-cultures with glioma cells under normoxic (CC) or hypoxic conditions (CC+Hyp) in our RNAseq study. Firstly, we found that most of the genes selected from the hypoxic-GAMs clusters (“hypoxia score”, **Supplementary Table S1**) were upregulated by hypoxia in BV2 cells (**Figure 3A**). Secondly, numerous genes defining specific clusters were also significantly affected by hypoxia in BV2 cells, including myeloid markers common for Mg-GAMs and Mo/Mφ-GAMs (**Figure 3B**). Furthermore, we found that markers of IFN, Transitory and Ribo-GAMs were predominantly downregulated by hypoxia, while phagocytic and chemotactic GAMs genes were induced by hypoxia in BV2 cells (**Figure 3C**). These findings are in agreement with previous reports, showing that ribosomal and RNA processing genes are downregulated in response to hypoxia ^33,34^. On the other hand, the chemotaxis and phagocytic phenotype genes upregulated in response to hypoxia suggest the active rearrangements in the cytoskeleton and increased motility of these myeloid cells facilitating cell migration and clearing of the necrotic debris ^3,8^. The genes from the lipid-GAMs cluster were mostly upregulated in hypoxic co-cultures, including *F10, Lgals3, Gpnmb, Apoe* or *Ctsb* (**Figure 3C**). However, some markers such as *Ftl1, Fth1* or *Fabp5* were found significantly downregulated, suggesting that the overall role of hypoxia in driving the lipid phenotype in TAMs is complex (**Figure 3C**). In many cases, the hypoxia in the BV2-glioma co-culture had a similar effect on the expression of the tested markers as hypoxia alone, although for some single genes, the differences were noted, as in the case of *Cd83* or *Cx3cr1* (**Figure 3B-C**).

**Figure 3.**
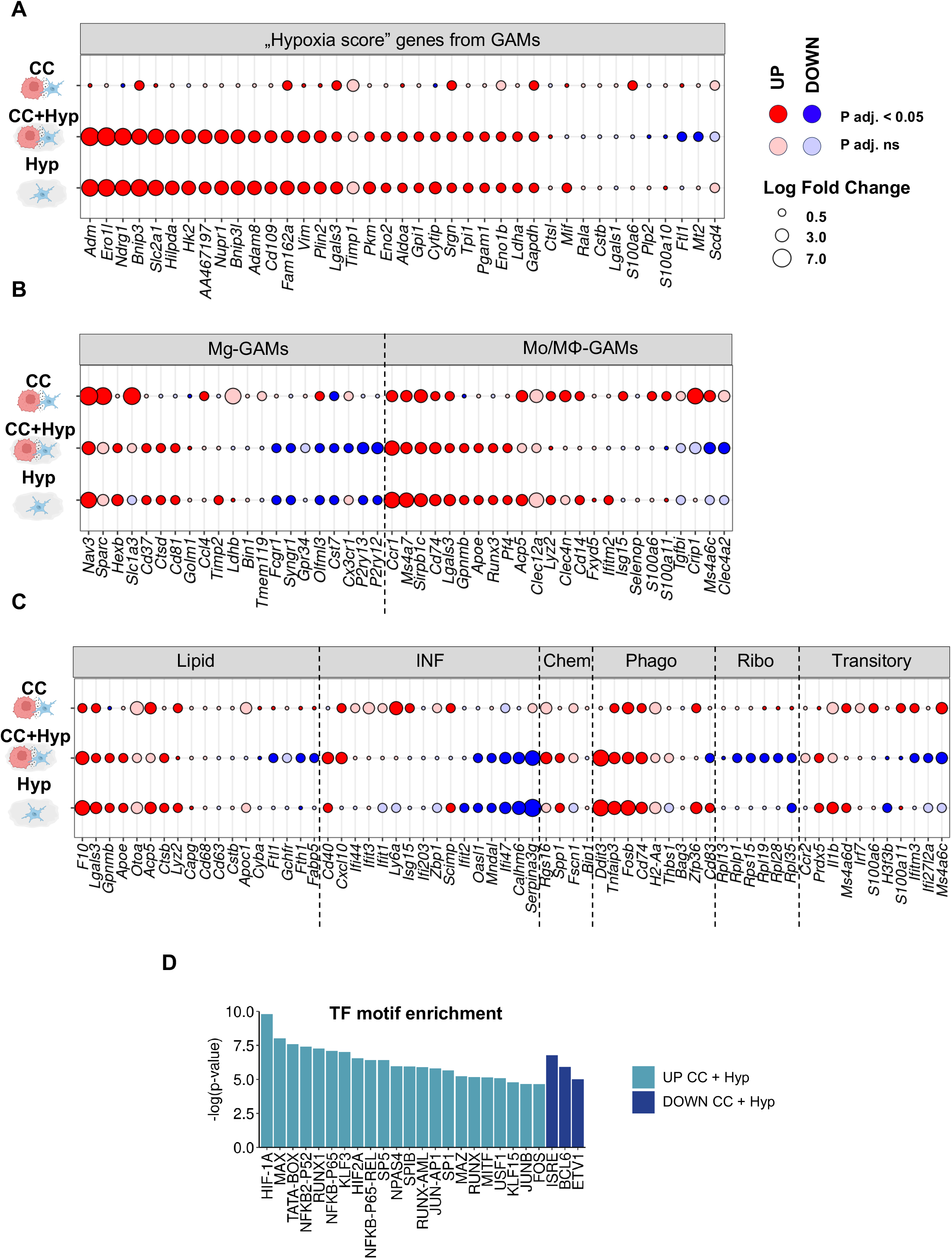
Hypoxia dysregulates the expression of multiple monocytic and microglial GAM subcluster marker genes. **A,** Gene expression changes in RNAseq data from Figure 2 in BV2 cells in response to co- culture with glioma GL261 cells in normoxia (CC), in response to co-culture with glioma GL261 cells in hypoxia (CC+Hyp) and in response to hypoxia only (Hyp) are shown for the genes included in “hypoxia score” for GAMs. The size of the circle indicates the fold change in a particular comparison. Dark blue and red colours indicate statistically significant gene expression changes (with p adjusted value < 0.05), while pale blue and pale red colours indicate non-significant regulation (with p adjusted value > 0.05). **B,** Gene expression changes in BV2 cells in comparisons as in (**A**) for top Mg-GAMs and Mo- GAM marker genes. **C,** Gene expression changes in BV2 cells in comparisons as in (**A**) for myeloid marker genes representing specific subpopulations of Mo-GAMs, including lipid, interferon (IFN), chemotactic (Chem), phagocytic (Phago), ribosomal (Ribo) or transitory monocytes. **D**, Transcription factor (TF) motif enrichment analysis on the promoters of significantly up- or down-regulated genes from (**A-C**) in BV2 cells after hypoxic co-culture with glioma cells.

To dissect regulatory networks, we searched for the top TF motifs that were enriched at the promoters of hypoxia-upregulated or downregulated myeloid marker genes analysed here. In addition to the enriched HIF-1/2A binding sites, other TF motifs were also detected in hypoxia-increased genes, including those for MAX, RUNX, NF-κB, AP1 and others (**Figure 3D**). Motifs for ISRE, BCL6 and ETV1 were found as enriched in the promoters of hypoxia-downregulated genes (**Figure 3D**). Overall, the results indicate that hypoxia has a dominant effect on the expression of numerous GAM markers, which in turn may affect the function of particular myeloid cells present within the TME, with a potential role in supporting tumorigenesis.

To study whether hypoxia-reprogrammed myeloid cells support glioma aggressiveness, we also performed an RNAseq analysis on GL261 glioma cells from monocultures and co-cultured with BV2 cells (**Figure 4A**). The PCA showed a very strong effect of hypoxia on sample separation either in co-culture or GL261 alone (**Figure 4B**) and numerous transcriptomic changes in GL261 cells in response to hypoxia in co-culture with BV2 or to hypoxia alone were identified (**Figure 4C-E**). Overall, the hypoxic stress induced a typical hypoxia response in both cases, as determined with the ORA GO analysis (upregulation of processes related to decreased oxygen levels, angiogenesis, etc.; **Figure 4F-G** and **Supplementary Table S4**). However, the most significantly enriched process within upregulated genes for glioma cells in co-cultures under hypoxia was a positive regulation of cell migration, while for hypoxic GL261 monocultures the top enriched pathways included fat cell differentiation and response to oxygen levels (**Figure 4F and G**, respectively and **Supplementary Table S4**). To test the role of hypoxia further, we used a Matrigel invasion assay, in which glioma cells were invading through the Matrigel-coated inserts towards the signal released by microglia cells present at the bottom of the well (**Figure 4H**). Indeed, the most efficient transmigration was observed for glioma cells exposed to both hypoxic stress and presence of microglia (**Figure 4I**). These results indicate that the presence of microglial cells under hypoxic conditions enhanced the migratory and invasive phenotype of glioma cells, which is in line with the more aggressive phenotype observed in patients with hypoxic tumours.

**Figure 4.**
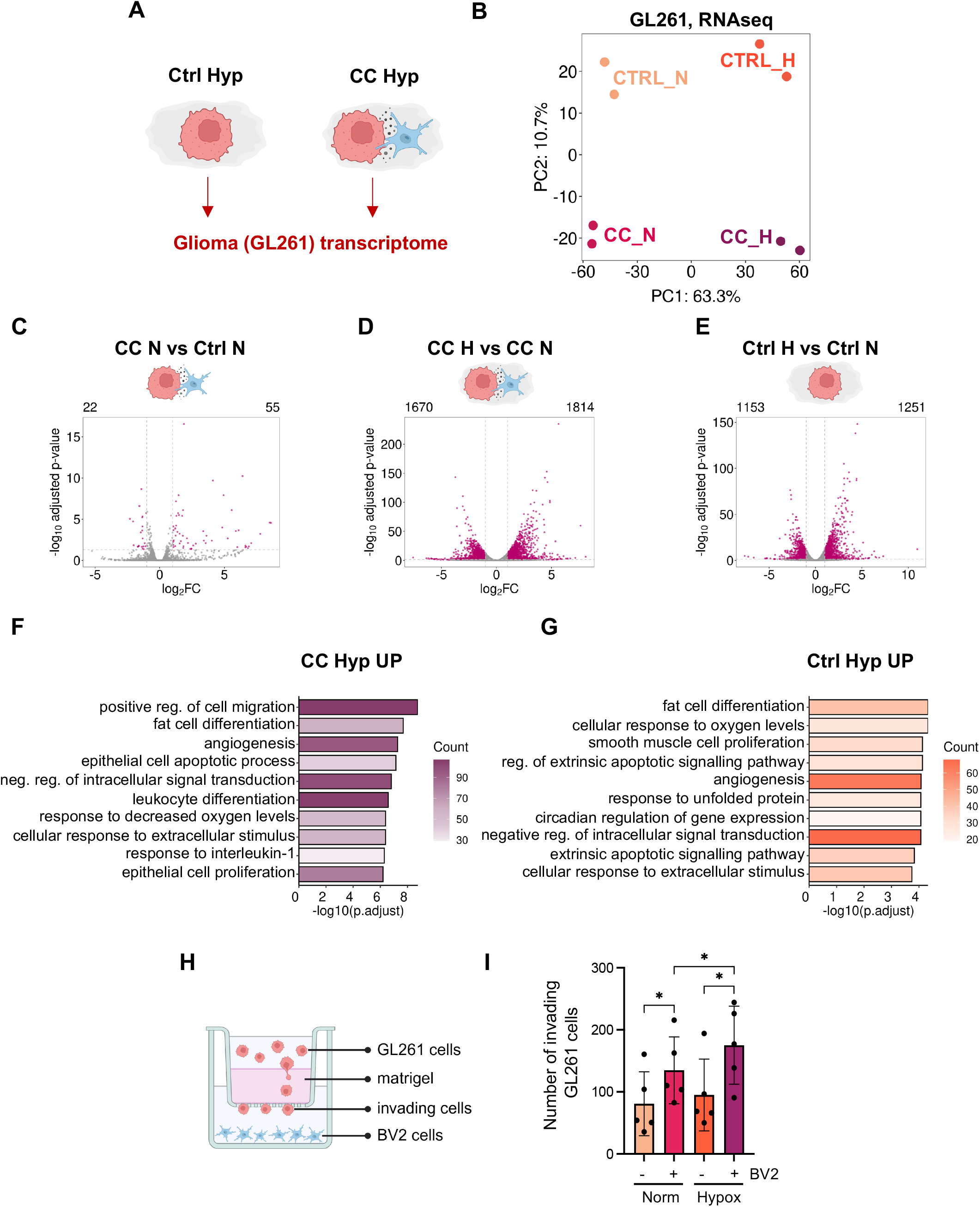
Hypoxia-dependent gene expression analysis in GL261 cell monocultures and co-cultures with microglia. **A,** A scheme indicating the analysis of GL261 transcriptome in co-culture with microglial cells or in monocultures. **B,** A principal component plot of normalised gene expression for GL261 cells in normoxia (Ctrl_N), in hypoxia (Ctrl_H) or in normoxic co-cultures with microglia (CC_N) and in hypoxic co-cultures with microglia (CC_H). **C-E,** Volcano plots showing differential gene expression in GL261 cells co-cultured with BV2 (**C**) in normoxia, (**D**) in hypoxia or (**E**) as hypoxic monocultures. **F-G**, Overrepresentation analysis of GO processes enriched in GL261 at the differentially up- regulated genes in hypoxic glioma co-cultures with microglia (**F**) or in hypoxic monocultures (**G**). **H-I,** A scheme representing Matrigel invasion assay. Glioma cell invasion in the absence or presence of microglial BV2 cells was measured after 16 h of co-culture in normoxia or hypoxia (<0.1% O2). A scheme showing invasion assay (**H**) and quantitation of invading GL261 cells (**I**). The mean with standard deviation (SD) is shown from five independent biological replicates. Statistical significance was assessed using one-tailed Wilcoxon matched-pairs test (* *p*<0.05).

### Glioma and hypoxia exposure reprograms the epigenome of BV2 cells

It was previously shown that hypoxia can significantly alter the chromatin landscape in cancer cells ^34–39^. Therefore, we considered hypoxia-induced chromatin remodelling as a causative factor for differential gene expression in microglia. First, we performed an immunofluorescent (IF) staining of key histone H3 methylations to verify whether hypoxia affects these marks in microglia co-cultured with glioma cells. As shown, hypoxia globally increased H3K27me3 or H3K9me3 marks both in glioma and in BV2 cells (**Supplementary** Figure 5A-D). This prompted us to investigate the chromatin accessibility patterns in glioma-interacting microglial cells in normoxic and hypoxic conditions using ATACseq assay (**Supplementary** Figure 5E). Again, to prevent any reoxygenation-dependent chromatin changes, we performed an ATACseq assay optimised for formaldehyde-fixed cells, prior to the separation of co-cultures with flow cytometry and further processing ^40^. Overall, we detected 26718 ATACseq peaks across all conditions, of which 0.1% were specific to Ctrl_N, 6.5% specific to CC_N, 14.5% specific to Ctrl_H and 11.3% specific to CC_H (**Supplementary** Figure 5F). Interestingly, the PCA on ATACseq peaks showed that co-culture with glioma cells had a much stronger effect on the chromatin accessibility in BV2 cells (separation along the PC1 axis) than hypoxia (separation along the PC2 axis) (**Figure 5A**). The co-culture of BV2 with glioma cells in normoxia resulted in a larger number of differentially increased than decreased chromatin accessibility peaks in BV2 cells (2173 up and 716 down peaks, respectively; **Supplementary** Figure 5G). However, the co-culture of BV2 with glioma cells in hypoxia further dysregulated chromatin accessibility in BV2 cells in comparison to glioma-co-culture in normoxia or to hypoxia-alone (4418 up and 3885 down peaks in hypoxic coculture; 1831 up and 1514 down peaks in hypoxia alone control; **Supplementary** Figure 5H-I). We then analysed the genomic localisation of differentially regulated peaks and found specific differences in the number of promoter-, intron- and distal intergenic-associated peaks in normoxic BV2 co-cultures or in hypoxic BV2 monocultures (differences in up versus down peaks in CC and in Hyp alone; **Figure 5B**). Namely, co-culture in normoxia resulted in the upregulation of promoter, intronic and distal intergenic peaks (**Figure 5B**, left panel), while hypoxia alone led to the downregulation of promoter peaks (**Figure 5B**, right panel). Importantly, a combination of these two factors (co-culture in hypoxia) resulted in a relatively similar number of up and downregulated peaks at particular genomic regions (**Figure 5B**, middle panel). Further analysis of the upregulated peaks, specifically at the promoter regions, showed that 109 peaks out of 1680 CC_Hyp UP peaks (6.49%) were also increased due to hypoxia alone, while 1515 upregulated peaks required specific concomitant exposure to hypoxia and glioma (**Figure 5C**). Amongst the downregulated promoter peaks (**Figure 5D**), 483 (33.56%) out of 1439 CC_Hyp DOWN peaks were also downregulated due to hypoxia alone, and 951 required specific concomitant exposure to hypoxia and glioma for their downregulation. This suggested that hypoxia as a single factor had a stronger impact on the downregulation rather than on the upregulation of promoter region openness (**Figure 5B-D**). In turn, the combined factors of glioma presence and hypoxia were important for gaining accessible promoters in hypoxic BV2 cells.

**Figure 5.**
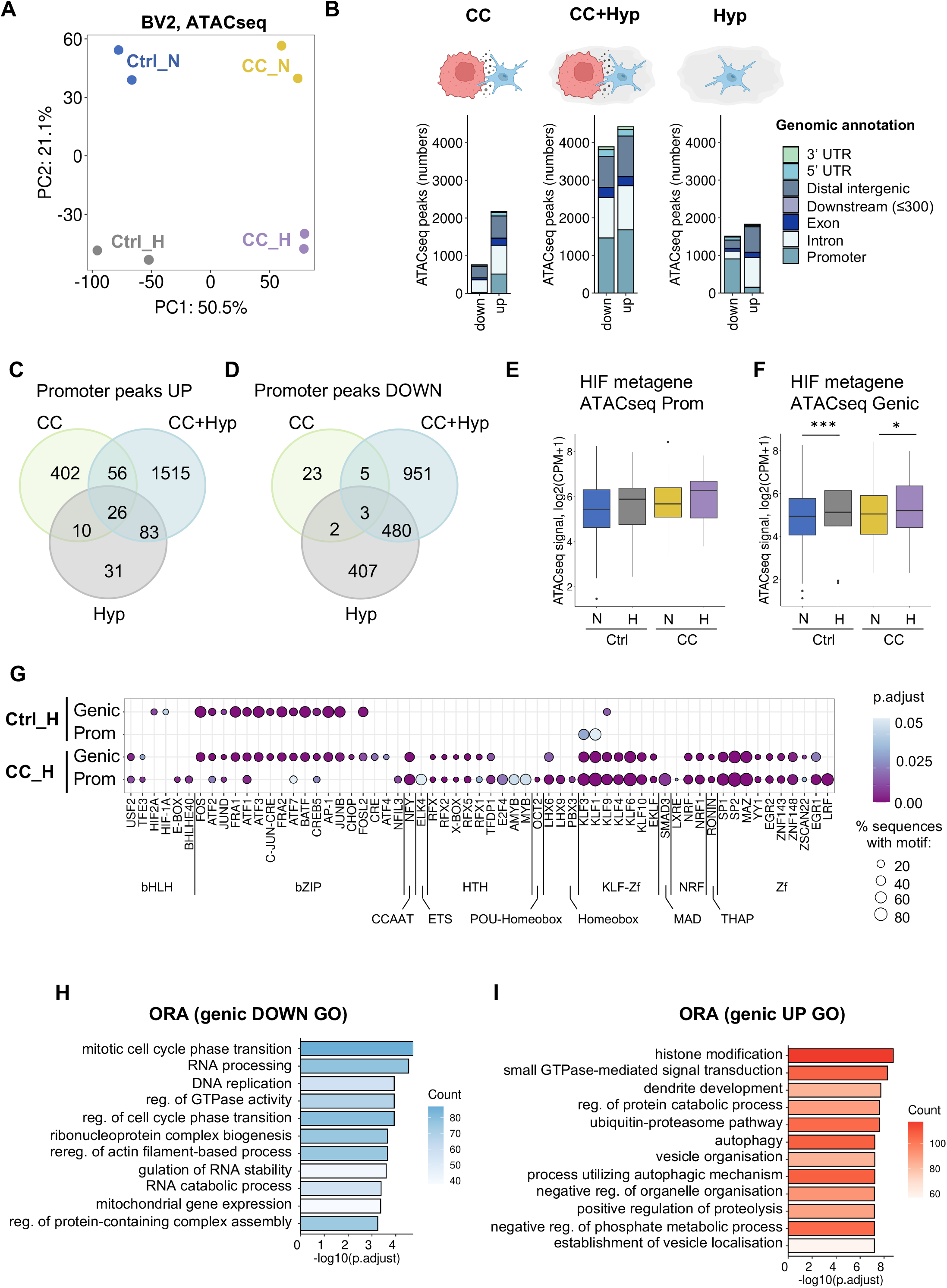
Glioma and hypoxia exposure reprograms the epigenome of BV2 cells. **A,** Principal component plot on identified ATACseq peaks in BV2 cells treated with hypoxia (<0.1% O2) or normoxia in the presence or absence of glioma cells. **B**, Bar plots showing differentially regulated ATACseq peak counts annotated to specific genomic regions, as indicated in the figure legend. Three differential analyses were performed for chromatin accessibility in BV2 cells: in response to co-culture with glioma GL261 cells in normoxia (CC; left graph), BV2 in response to co-culture with glioma GL261 cells in hypoxia (CC+Hyp; middle graph) and in response to hypoxia only (Hyp; right graph). **C-D**, Venn diagrams summarising common and distinct upregulated (**C**) or downregulated (**D**) ATACseq peaks at promoters in BV2 cells treated with CC, CC+Hyp or Hyp. **E-F,** Analysis of HIF metagene score in ATACseq pekas at the **(E)** gene promoter and **(F)** promoters and gene bodies (denoted as genic) in BV2 cell in hypoxia. HIF metagene set was defined by Lombardi et al ^26^. The average ATACseq signal of the HIF metagene score is represented as read counts per million. Two-way ANOVA and post-hoc Tukey test determined the statistical significance. **G,** Enrichment of TF motifs in ATACseq peaks at the gene promoter or genic regions that increased in BV2 cells treated with hypoxia in co-culture with GL261 (CC) or alone (Ctrl) in comparison to respective CC or Ctrl normoxic cells. Circle size indicates the percentage of sequences in the subset with the motif, while the colour gradient reflects the significance of enrichment. All indicated enriched TF motifs are statistically significant (p adjusted<0.05). At the bottom part of the graph, the TFs are additionally grouped into respective TF families: bHLH - basic Helix-Loop-Helix; bZIP - basic Leucine Zipper; CCAAT - TF that specifically bind to the CCAAT box; ETS - E26 Transformation-Specific; HTH - Helix-Turn-Helix; POU- Homeobox - TFs that contain both POU-specific domain and a homeodomain; Homeobox - TF that contain homeodomain; KLF-Zf - Krüppel-Like Factors, Zinc Finger; MAD – ‘mothers against decapentaplegic’; NRF - Nuclear Respiratory Factor; THAP - Thanatos Associated Proteins; Zf - Zinc Finger. **H-I,** Overrepresentation analysis showing GO processes in downregulated (**H**) or upregulated (**I**) genic ATACseq peaks. The top 15 GO processes were selected based on the lowest adjusted *p*-values, and closely related terms were merged using REVIGO. The gradient of colours indicates the gene count in each associated process.

Further analysis of promoter peaks and genic peaks (including the promoter and gene body) assigned to particular genes showed that the ATACseq peaks allocated solely to the promoters of the HIF-inducible genes were not significantly increased in response to hypoxia (**Figure 5E**) ^26^. However, including genic peaks in the analysis of the HIF signature showed significant upregulation of ATACseq peaks in the HIF-inducible genes in response to hypoxia (and more significant in control hypoxia samples), suggesting that peaks outside of the promoters were particularly induced in response to hypoxia at the HIF-target genes (**Figure 5F**) ^26^. This result was additionally confirmed by the TF enrichment analysis on all upregulated ATACseq peaks, where HIF-1A and HIF-2A enrichment was only found in genic, but not promoter-only regions in hypoxic cells from monocultures (**Figure 5G**).

In addition, a number of TFs were enriched at the genic peaks both in co-cultured and control hypoxic BV2 cells, including AP1 (FOS and JUN subunits), ATF1-4, FRA1-2, CHOP, and others (**Figure 5G**). TF motifs that were the most specifically and significantly enriched in upregulated peaks in hypoxic co-cultures included the ones that have been implicated in the regulation of microglial cell functions (**Figure 5G**). For example, we found motifs for KLF6 and EGR1, which play a role in the regulation of microglial activation and inflammatory responses ^41,42^, for KLF4 which is involved in microglial cell polarization and differentiation ^43^, for KLF9 known to regulate oxidative stress responses in microglia ^44^, for SP1 involved in microglial cell activation, survival and proliferation ^45^ and SMAD3 regulating activity, differentiation, and maintaining a quiescent state of microglial cells ^46^. Similarly, as in the RNAseq analysis, the ORA GO analysis at decreased genic ATACseq peaks showed downregulation of processes related to mitotic cell division, GTPase activity, DNA replication and various RNA processing mechanisms (**Figure 5H** and **Supplementary Table S5**). On the contrary, the upregulated genic ATACseq peaks were associated with histone modification, dendrite development, autophagy, vesicle organisation and other processes (**Figure 5I** and **Supplementary Table S5**).

Our data so far suggested that the glioma co-culture has the strongest influence on the chromatin accessibility changes in microglia specifically in hypoxic conditions (**Figure 5B**). Therefore, we verified which peaks and associated processes were the most enriched during glioma co-culture in hypoxia when compared to hypoxic monocultures. Out of 4418 peaks upregulated in co-cultured hypoxic BV2, 2257 peaks were more enhanced in hypoxia co-culture in comparison to hypoxic BV2 monocultures (**Supplementary** Figure 6A). Out of 3385 downregulated peaks in co-cultured hypoxic BV2, 1638 were more decreased in hypoxia co-culture in comparison to hypoxic BV2 monocultures. The genomic annotation analysis of these peaks suggested that the combined effect of hypoxia and glioma co-cultures when compared with hypoxia-alone led to an increase in chromatin accessibility at the promoter regions and a decrease in accessible introns and distal intergenic chromatin sites in BV2 cells (**Supplementary** Figure 6B). The ORA GO analysis of these enhanced changes in promoters with increased accessibility specifically in co-cultured hypoxic BV2 cells showed upregulation of processes such as autophagy, histone modifications, cellular components disassembly etc (**Supplementary** Figure 6C-D). Interestingly, while there was a low number of genes with the decreased chromatin accessibility at the gene promoters in response to co-culture and hypoxia, these genes were associated with the neuroinflammatory responses, defence responses or leukocyte activation (**Supplementary** Figure 6C-D). This suggests that hypoxia and glioma exposure by reducing chromatin accessibility supports the immunosuppressive phenotype of BV2 cells. In turn, processes associated with the changes at the intronic and exon peaks specifically in hypoxic co-cultures were associated with tissue specific differentiation processes in upregulated peaks and regulation of cellular signalling, including GTPase activity, calcium ion homeostasis, metal ion transport etc in downregulated peaks, respectively (**Supplementary** Figure 6E).

### Concordant hypoxia-dependent transcriptomic and epigenomic changes in glioma-co-cultured BV2 cells impact myeloid cell functional markers

Subsequently, we compared differential changes in the gene expression and chromatin accessibility in the glioma-co-cultured BV2 cells. The Spearman and Pearson correlation coefficients (0.24) for ATACseq peaks and gene expression indicated a partial but significant correlation of affected genes in both datasets (**Figure 6A**). Out of 2605 genes with induced expression, 645 had concordant changes in the chromatin accessibility (24.76%) (**Figure 6B**). For the 1942 genes with decreased expression, 345 had concordant downregulation of chromatin accessibility in the same condition (17.76%) (**Figure 6C**). The ORA GO analysis on the concordant upregulated genes indicated response to hypoxia as the top pathway, but also other quite diverse biological processes, including regulation of transcription in response to stress, neuron migration, muscle organ development and other (**Supplementary** Figure 7A and **Supplementary Table S6**). The concordant downregulated again genes indicated typical processes known to be downregulated in hypoxia, including ribosome biogenesis, DNA replication, cell cycle division etc (**Supplementary** Figure 7B and **Supplementary Table S6**). Very few of the concordant changes were additionally enhanced in BV2 cells co-cultured with glioma in hypoxia when compared to hypoxia alone (29 genes up and 10 genes down). However, among the genes specifically downregulated in hypoxic co-cultures there were genes falling into significantly enriched GO processes, such as semaphorin-plexin signalling pathway involved in neuron projection guidance, signal release, regulation of secretion by cell, but also again myeloid leukocyte differentiation (**Supplementary** Figure 7C and **Supplementary Table S6**). These processes represent homeostatic functions of microglia ^3,5^.

**Figure 6.**
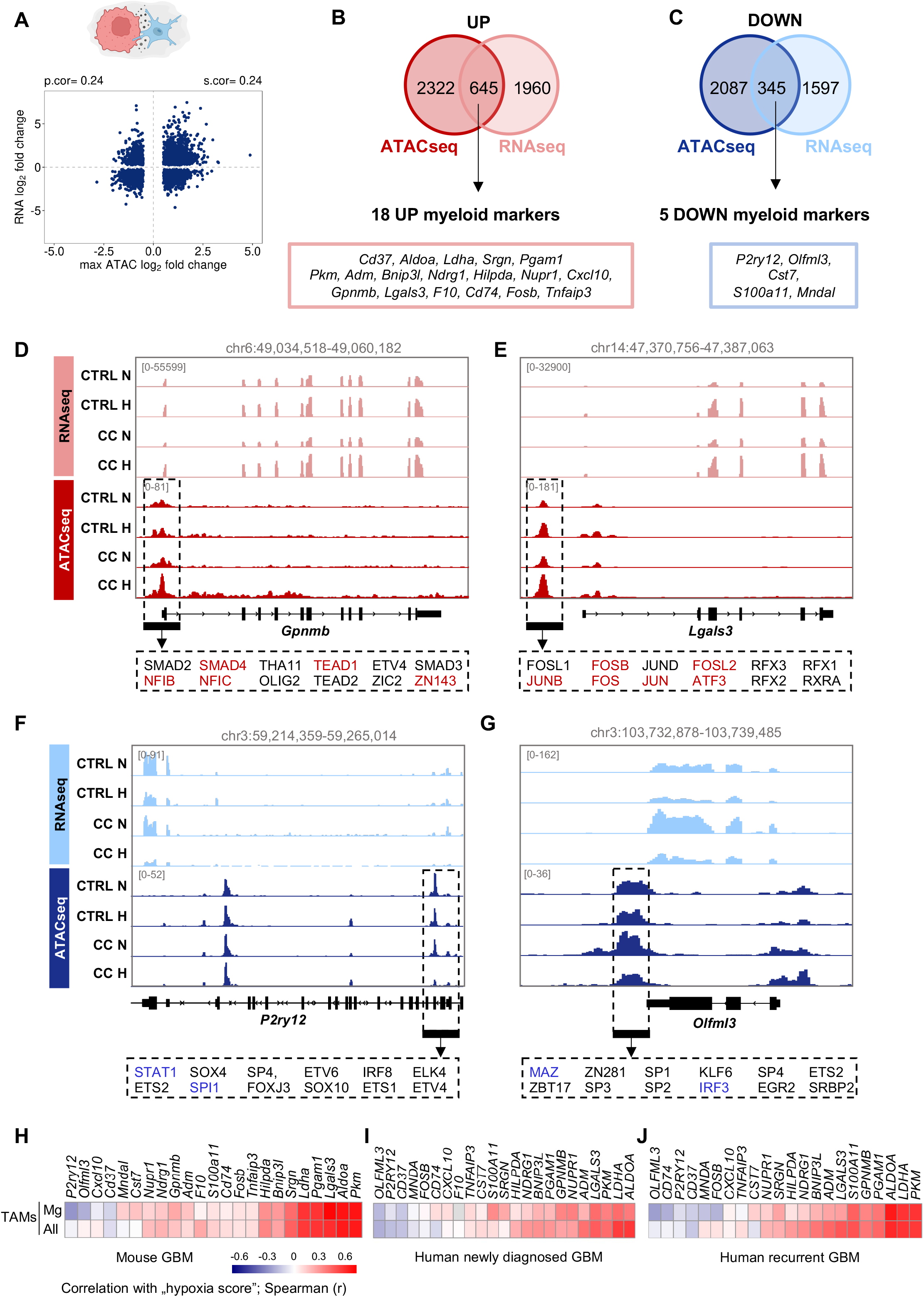
Concordant hypoxia-dependent transcriptomic and epigenomic changes in glioma-co-cultured BV2 cells impact myeloid cell functional markers. **A**, A dot plot showing the correlation of mRNA expression changes versus maximal ATACseq peak fold changes per gene. Statistically significant pearson (p.cor) and spearman (s.cor) correlation coefficient values are shown. **B-C**, Venn diagrams showing concordantly upregulated (**B**) or downregulated (**C**) gene expression and corresponding genic ATACseq peaks in BV2 cells treated with hypoxic co-culture with glioma cells. The rectangular frames indicate myeloid marker genes from Figure 3 identified in the concordant ATACseq and RNAseq changes. **D-G,** The mRNA expression (RNAseq) and ATACseq profiles (from Integrative Genomics Viewer, IGV) for concordantly upregulated (*Gpnmb* (**D**) and *Lgals3* (**E**)) and downregulated (*P2ry12* (**F**) and *Olfml3* (**G**)) genes in BV2 cells treated with hypoxia co-culture with GL261 cells. The FIMO analysis was used for the selected ATACseq peaks, to identify available motifs for TFs, as indicated in rectangular boxes under each gene. The IGV track shown are the average from two biological repeats. **H-J**, Heatmaps representing the correlation matrix between hypoxia score and concordantly hypoxia-regulated genes at the expression and chromatin level indicated in (**B**) and (**C**) in scRNAseq datasets for mouse TAMs (**H**), TAMs in human newly diagnosed GBM (**I**) and TAMs in human recurrent GBM (**J**) (all from GSE163120) ^6^.

The downregulation of the myeloid leukocyte differentiation process in concordant changes brought us to our initial findings that some myeloid markers or GAM cluster genes might be dysregulated in hypoxia through chromatin remodelling. We found that out of 53 transcriptionally induced GAM-associated marker genes (listed in **Figure 3**), 18 (34%) had concordant changes in the chromatin accessibility in BV2 cells co-cultured with glioma in hypoxic conditions (**Figure 6B**, lower box). Similarly, for the significantly downregulated GAM-associated marker genes, 5 out of 32 (15.6%) were concordantly decreased at the mRNA and chromatin accessibility level (**Figure 6C**, lower box). Of the concordantly dysregulated myeloid cell markers in hypoxia, some were already described as crucial for the tumour-supportive role and lipid metabolism in GAMs or directly in immunosuppression. *Gpnmb* was recently shown to play an immunosuppressive role in the tumour microenvironment of glioma as its deletion increased the inflammation and reduced the tumour size ^47^. We have scanned for the presence of specific TF motifs in the upregulated chromatin accessibility peak in the *Gpnmb* promoter and found SMAD2, SMAD4, NFIB and NFIC among the most frequent TF motifs occurring within this peak (**Figure 6D** and **Supplementary Table S7**). The TGFB/SMAD axis is already known to regulate the immunosuppressive phenotypes in myeloid cells ^48^. In turn, at the *Lgals3* promoter, the upregulated accessible chromatin region was abundant with binding motifs for AP-1 and ATF3 TFs (**Figure 6E** and **Supplementary Table S7**), which can mediate the tumour-supportive function of macrophages, e.g. via upregulation of lipid metabolism ^49^. Moreover, both AP1 and AFT3 are known to be active in hypoxic conditions and may well play a significant role in the regulation of functions of hypoxic microglia ^29^. Interestingly, for the downregulated genes, at the *P2ry12* Mg-GAM marker gene promoter the ATACseq peak was almost completely lost in BV2 cells after co-culture with glioma cells in hypoxic conditions (**Figure 6F** and **Supplementary Table S7**). The TFs with binding motifs in this peak included SPI1 (also known as PU.1) and IRF8, both known to play essential roles in the regulation of microglial cell identity and function, as well as regulation of *P2ry12* expression (**Figure 6F** and **Supplementary Table S7**) ^50–52^. In addition, we found that expression of *Spi1* decreased in hypoxic conditions in our RNAseq analysis, which may additionally contribute to the downregulation of *P2ry12* expression in hypoxia. Motifs for other TFs involved in myeloid cell functions were also identified in this peak, including STAT1 and ETS1/2, although these have not been previously directly linked to the regulation of *P2ry12* expression ^53^. Regulation of expression of *Olfml3* in BV2 cells exhibited an interesting pattern, as it clearly followed the changes in the chromatin accessibility at its 3’UTR region (**Figure 6G**). Co-culture with glioma cells upregulated the expression of *Olfml3* in BV2 cells, which was then inhibited in hypoxic conditions. From the TF motifs found in the hypoxia-regulated chromatin accessibility peak in the *Olflm3* gene, the IRF3, SP1, and ETS2 TFs were previously linked with the regulation of immune responses in myeloid cells (**Figure 6G** and **Supplementary Table S7**) ^45,54,55^. Overall, our data indicate that although chromatin accessibility changes only partially correspond with the strong gene expression changes in hypoxia, some of these concordant alterations may well play a role in defining the myeloid cell properties.

Finally, we tested if the myeloid markers genes that were concordantly regulated at the chromatin and mRNA levels in hypoxic co-cultures, correlate with the “hypoxic score” in GBM GAMs *in vivo*. The correlation analysis in mouse and human GBM Antunes-datasets between the “hypoxic score” and expression of particular genes indicated *P2ry12* and *Olfml3* as negatively correlated and *Gpnmb*, *Lgals3*, *Tnfaip3*, *Cxcl10*, as well as hypoxic TAM markers as positively correlated either in all TAMs or at least in Mg-TAMs (**Figure 6H-J** and **Supplementary Table S8**). Interestingly, some of the genes were inversely correlated with “hypoxic score” than determined in the co-culture model, including *Mndal*, *Cd37* or *S100a11*.

We cannot exclude that other factors could contribute to their expression in hypoxic cells *in vivo*.

## DISCUSSION

The role of hypoxic stress as a factor affecting myeloid cell identity has not been addressed up to date. Using a well-controllable *in vitro* model of the direct glioma-microglia co-culture, we show that hypoxia is a strong factor significantly reprograming the chromatin accessibility and gene expression in microglial cells that are exposed to glioma cells. The analysis of concordant changes in chromatin accessibility and gene expression indicates upregulation of genes associated with tumour-supportive functions of microglia, while genes involved in homeostatic and neuroprotective functions, along with canonical microglial markers, are downregulated. We confirm that many of hypoxia-induced changes identified in this *in vitro* model can be observed in human and mouse GBM mRNA samples *in vivo*. We provide the evidence that most of the functional subpopulations of GAMs may be influenced to some extent by internal hypoxic tumour stress and a more detailed analysis of microglial, monocytic and macrophage clusters in single-cell RNAseq data might be required to dissect these distinct hypoxia-dependent changes.

The “hypoxia score” in GAMs, which reflects the influence of hypoxic stress, often correlates with the expression of functional lipid GAM marker genes, e.g. *Lgals3* or *Gpnmb*, which is in line with studies showing that hypoxia regulates lipid metabolism, lipid processing and storage ^56^. However, some of the lipid markers in GAMs, in addition to lipid uptake and metabolism, can also be secreted and play versatile roles in tumours. For example, Galectin-3 *(Lgals3*) expressed in GAMs was already shown to stimulate glioma cell migration and invasion, impair the inflammatory phenotype and overall promote the immunosuppressive environment ^57^. In breast cancer, the expression and secretion of Galectin-3 in TAMs was shown to be induced in hypoxic conditions, and again, had a pro-tumorigenic role. The authors showed that inhibition of Galectin-3 significantly improves the effect of anti-angiogenic therapy in breast cancer model ^58^. Although studies to date indicate that *Lgals3* expression is already much higher in Mo/Mφ-GAMs, which are more abundant in the tumour core, than in Mg-GAMs ^5,6^, the further upregulation of *Lgals3* in all hypoxic GAMs and overall in all cells present in hypoxic regions, including glioma cells, may have a cumulative effect in supporting the glioma progression. Further studies are needed to explore the therapeutic potential of targeting Galectin-3 in GBM, particularly in tumours with high hypoxia levels.

We show that tumour hypoxia-dependent reprogramming of chromatin can act as a master regulator of some changes that are essential for the myeloid cell identity. The TFs IRF8 and SPI1 are known to be crucial for the regulation of identity and specification of microglial cells. ^50–52^. We observe that reduced expression of microglial marker *P2ry12* in hypoxia is associated with downregulation of chromatin accessibility at the promoter region of this gene, which also contains the binding sites for IRF8 and SPI1. Moreover, such downregulation of chromatin accessibility and gene expression of *P2ry12* is only observed when microglia cells are exposed to hypoxia in the presence of glioma cells and not when microglia are grown as monocultures. This suggests that multiple stimuli are required, i.e. contacts with glioma cells and loss of oxygen, which synergise in decreasing the chromatin accessibility at the *P2ry12* promoter. Further studies are needed to dissect the mechanism of such co-regulation in GAMs.

We report that hypoxia impacts expression of the top GAM genes that are used as markers to distinguish between the tumour-infiltrating microglia and peripheral monocytes/macrophages, including *P2ry12* and *Lgals3* genes, respectively. As many studies rely on those markers in dissecting myeloid subpopulations in tumours, the awareness that their expression might be affected by hypoxia may help to resolve many conflicting data.

While hypoxia imposes extensive global changes in the chromatin accessibility in BV2 cells, not all of these changes are concordant with the gene expression in these cells. Several scenarios could explain these discrepancies. It is possible that the chromatin accessibility-dependent changes require more time under hypoxia to translate into actual gene expression changes, while the HIF family of TFs or other TF-mediated transcriptional changes are more rapid and direct, allowing a very quick adaptation to hypoxic conditions ^59^. Moreover, some chromatin accessibility alterations in hypoxia might be the result of changes of 3D contacts in the chromatin structure and do not necessarily directly influence the expression of nearby genes ^60^. Therefore, deeper epigenetic studies are needed, which capture chromatin conformation in hypoxia, to further examine the significance of all chromatin accessibility changes in glioma- co-cultured hypoxic microglial cells.

While the *in vitro* direct co-culture system of microglial cells with glioma cells with or without hypoxia has some limitations, it reliably reflects chromatin openness and transcriptomic changes underlying reprograming of GAMs, as similar patterns are detected in published scRNAseq *in vivo* data. While it is a simplified model and does not incorporate other cells from the TME, it allows to dissect discrete changes occurring at the beginning of the reprograming both in glioma and microglial cells and to unravel the impact of hypoxia. Some key features of the pro-tumoral reprogramming of GAMs by hypoxia detected at the transcriptomic and epigenomic levels in our co-culture model we were able to confirm through the morphologic transformation and increased invasion-promoting activity of microglia upon exposure to hypoxia and contacts with glioma cells.

## CONCLUSIONS

Altogether, our data shows that hypoxia has a strong impact on the reprogramming of myeloid cells and expression of markers, which define the cell identity and/or phenotypic state. To a large extent, the changes in gene expression are likely to be regulated through the immediate action of TFs activated in hypoxia, including HIF-1α/HIF-2α, NF-κB, RUNX1, AP1 etc. However, in some cases, hypoxia-dependent reprogramming of chromatin will also regulate the expression of myeloid marker genes, as shown for *P2ry12* or *Olfml3*-microglial markers or lipid-GAMs related genes such as *Lgals3* or *Gpnmb*. Further studies are warranted to test the clinical potential of targeting hypoxia-dependent epigenetic and transcriptomic changes in GAMs. Here, we provide a robust and reproducible model for studying the direct glioma-microglia interactions, which involves cross-linking-preserved oxygen-regulated changes and is suitable for the dissection of hypoxia-dependent mechanisms in GAMs and glioma cells.

## MATERIALS AND METHODS

### Cell lines

GL261 mouse glioma cells stably expressing pEGFP-N1 were cultured in DMEM with 10% FBS) and were obtained from Prof. Helmut Kettenman (MDC, Berlin, Germany) ^5,24^. The murine immortalised BV2 microglia cell line (received from Prof. Klaus Reymann from Leibniz Institute for Neurobiology) and the murine macrophage cell line RAW 264.7 (from Biological Resource Center ICLC Cell bank, #ICLC ATL02001) were cultured in DMEM with GlutaMAX™, supplemented with 2% and 10% FBS, respectively. Bone marrow-derived macrophages (BMDMs) were obtained from C57BL/6 mice at the age of 10-16 weeks, according to the protocol by Toda et al., with some modifications ^61^. In brief, bone marrow cells from the femur and tibia of mice were collected by a short centrifugation of the bones after removing the epiphyses. Erythrocytes were removed with the use of a dedicated lysis buffer (BD Pharm Lyse™ Lysing Buffer # 555899). Bone marrow cells were frozen in 10% DMSO/FBS. For differentiation into BMDMs, bone marrow cells were plated on glass coverslips in 6-well plates (1,6 mln cells/well) in DMEM w/Glutamax (Gibco 31966047) supplemented with 10% FBS and 50 ng/ml M-CSF. Culture medium was replaced with fresh medium (containing 10% FBS and 50 ng/ml M-CSF) at day 4 and 6. Additionally, M-CSF was added (50 ng/ml) to the medium on other days if the cultures were detaching from the plates. The BMDMs were used for hypoxic treatment on day 10, at which time the expression of macrophage markers was confirmed with qPCR (not shown). Antibiotics (100 U/mL penicillin, 100 µg/mL streptomycin) were supplemented in all cell culture media. All cell lines were maintained in a humidified atmosphere of CO2/air (5%/95%) at 37°C. The regular mycoplasma tests were carried out in all cells used.

### Hypoxic conditions

To create hypoxic conditions cells were incubated at <0.1 or 1% O2 in a M35 Hypoxystation (Don Whitley) hypoxic chamber in the presence of 5% CO2 at 37°C. The hypoxia-mimetic agent CoCl₂ (250 μM) was applied under standard culture conditions (21% O2).

### Direct co-culture assay

GL261 cells were seeded in DMEM supplemented with 10% FBS and antibiotics. After 24 h, the medium was changed to a microglia culture medium DMEM with GlutaMAX™ supplemented with 2% FBS and antibiotics and the BV2 cells were co-seeded onto GL261 cells at a ratio of 1:2 (BV2:GL261). Following 24h, the cells were cultured under normoxic or hypoxic conditions for another 16 hours and used in further assays. The control BV2 or GL261 cells were grown as monocultures end exposed to hypoxia or normoxia for 16 hours.

### Immunofluorescence

Cells were seeded on glass coverslips in 24-well plates in the same way as described in direct co-culture section. Cells were washed with PBS and 4% PFA was added to fix cells in hypoxic or normoxic conditions. After 10 min cells were washed with PBS followed by a 10-minute permeabilisation with ice-cold 100% methanol and a subsequent wash with PBS. After 1 hour blocking (3% donkey serum, 1% BSA, 0.3% Triton X-100, PBS) the H3K9me3 antibody (1:500, Millipore, Cat#07-442) or H3K27me3 antibody (1:500, Millipore, Cat#07-449) diluted in a blocking solution were added and incubated for 1 hour. This was followed by a triple wash in PBST and an hour incubation in the blocking solution with a secondary donkey anti-rabbit antibody conjugated with Alexa Fluor-555 (1:2000, Invitrogen, Cat#A31572,). Coverslips were washed 3x with PBST and stained with DAPI (1:1000) for 10 min. This was followed by a double wash with PBS and with distilled water and mounting on slides using a mounting solution (Dako).

### qPCR

RNA was isolated using RNeasy Mini Kit (QIAGEN), followed by quality and quantity assessment using NanoDrop 2000 Spectrophotometer (ThermoFisher). cDNA was synthesized from total RNA using SuperScript IV Reverse Transcriptase (Invitrogen). qPCR was performed with the SYBR Green PCR Master Mix Kit (Applied Biosystems) using QuantStudio 12K Flex Real-Time PCR thermocycler (Applied Biosystems) with a listed set of primers (**Supplementary Information**). The amplified product was normalized to the *Rn18s* housekeeping gene. The mRNA expression (fold change) was calculated using the 2–ΔΔCt method. The qPCR graphs show the mean ± SD of at least three biological replicates and statistical significance assessed with two-tailed student t-tests for each gene, performed with Graph Pad Prism software. Statistically significant *P*-value was considered below 0.05, with further details: * for *P*<0.05, ** for *P*<0.01 and *** for *P*<0.001.

### Western blotting

Cells were washed in PBS and lysed in SDS lysis buffer (10 mM Tris-Cl, pH 7.5, 0.1 mM EDTA, 0.1 mM EGTA, 0.5% SDS, 0.1 mM β-mercaptoethanol, protease/phosphatase inhibitors), followed by sonication and centrifugation, as described recently ^62^. Lysates with equal amounts of proteins were subjected to SDS-PAGE and western blotting. After blocking, the following primary antibodies were used: LGALS3 (1:1000, Biolegend. Cat#125401), HIF-1α (1:1000, Abcam, Cat#179483), β-actin (Sigma-Aldrich, Cat#A3854). An enhanced chemiluminescence detection system (ECL) and Chemidoc Imaging System (BioRad) were used to develop the signal from HRP-conjugated secondary antibodies.

### Invasion assay

The invasion assay was conducted as previously described ^63^. Briefly, BV2 cells (4×10⁴) were plated onto 24-well plates. The following day, GL261 cells (1×10⁵) were seeded into tissue culture inserts with Matrigel-coated membranes in a serum-reduced medium (2% FBS). After 24h of co-culture under normoxic (21% O2) or hypoxic (<0.1% O2) conditions, the invading GL261 cells were fixed, and their nuclei were stained with DAPI. The membranes from the inserts were excised and imaged using a fluorescence microscope (Leica DM4000B). The number of invaded cells across the entire membrane was counted. Each biological replicate (n=5) was performed in technical duplicate-wells.

### IvyGAP human glioblastoma data

Expression of hypoxia-inducible genes (*SLC2A1, VEGFA*), monocytic marker (*LGALS3*) or microglia markers (*P2RY12, TMEM119*) was tested in the transcriptional atlas of human glioblastoma, where samples for the RNAseq analysis had been collected via microdissection from anatomic structures within glioblastoma (Ivy Glioblastoma Atlas Project; http://glioblastoma.alleninstitute.org/) ^23^. The Z-score values were downloaded for the above genes. The violin plots showing the expression z-scores were generated with R programming environment (version 4.2.2, https://www.r-project.org/).

### Analysis of human and mouse scRNAseq data

UMAP plots were created using single-cell RNA sequencing data from mouse tumour-associated macrophages and two human tumour-associated macrophage datasets from Antunes et al., (all available at GSE163120) ^6^. The data were analysed in RStudio (R version 4.2.2 (2022-10-31); Platform: x86_64-pc-linux-gnu (64- bit); Running under: Ubuntu 22.04.1 LTS); BiocManager_1.30.19) and Seurat (v 5.0.1). UMAP dimensions and cell labelling were applied as in the original publication ^6^. Visualizations were generated using built-in Seurat functions, ggplot2 (v 3.5.1), Nebulosa (v 1.8.0)^64^, scCustomize (v 1.1.3) (https://doi.org/10.5281/zenodo.5706431), and RColorBrewer (v 1.1-3). Module “hypoxia score” (**Supplementary Table S1**) was computed using the Addmodulescore() function from the Seurat library. Correlations between selected marker genes and “hypoxia score” were calculated using the Spearman correlation coefficient with the cor() function. The values were visualized on heatmaps (pheatmap_1.0.12).

### RNAseq

Cells from hypoxia or normoxia conditions were fixed on ice with 3% Glyoxal fixation solution as described previously ^25^. For staining of microglia cells CD45-PE (1:400, Clone 30-F11; BD Pharmingen, Cat# 553081) was used. CD45+ cells and GFP+ cells were separated and collected using Cell Sorter BD FACSAriaII. RNA was isolated using RNeasy Plus Kit (Qiagen) according to manufacturer’s instructions and the quality was assessed with the Agilent 2100 Bioanalyzer with an RNA 6000 Pico Kit (Agilent Technologies). PolyA-enriched RNA libraries were prepared with the KAPA Stranded mRNA Sample Preparation Kit (Kapa Biosystems). The RNAseq libraries were sequenced at 2x151 bp paired-end on NovaSeq 6000 at the Laboratory of Sequencing (Nencki Institute). Two biological replicates per condition were analysed.

### RNAseq data processing and analysis

The quality of raw fastq data was assessed using FASTQC software (https://www.bioinformatics.babraham.ac.uk/projects/fastqc/). Low-quality reads and adapters were trimmed using Trimmomatic (version 0.36) ^65^. The resulting RNA sequencing reads were mapped to mm10 reference genome sequence using STAR aligner (version 2.6) ^66^. Read duplicates were marked with Picard Tools (version 2.17.1, https://broadinstitute.github.io/picard/) and only uniquely mapped reads were kept for the downstream analysis with samtools view -q 255. Quantification of mapped reads and summarisation by gene was performed using HTSeq-count (version 0.9.1), with overlap mode set to *union* and stranded mode to *reverse* (-m union -s reverse) ^67^. Differential analysis was performed using Bioconductor package DESeq2 ^68^. Genes that had significant (Benjamini and Hochberg-corrected P-value <0.05 and |log2 fold change| >=1) changes in their expression levels were called as differentially expressed. Gene ontology (GO) enrichment analysis was done using enrichGO form clusterProfiler ^69^. Revigo and Cytoscape (version 3.10.2, https://cytoscape.org/) were used to summarise and visualise the GO-enriched terms ^70^. Motif enrichment analysis for selected genes was performed using HOMER findMotifs.pl (http://homer.ucsd.edu/homer/motif/) for known TF motifs with the default parameters. All sequencing track were generated by Integrative Genome Viewer (IGV) using a normalized bigWig file created with bamCoverage of DeepTools and normalized to genome coverage - RPGC ^71,72^. The averaged signal across all regions for the two replicates in each condition was calculated using bigwigCompare of DeepTools ^71,72^ to generate a combined BigWig file. The RNAseq data generated in this project is deposited at the NCBI platform under GSE279536 accession number.

### ATACseq

ATACseq was performed as previously described with some modifications ^34^. While still in hypoxic conditions, cells were fixed with 1% formaldehyde (Thermo Scientific) for 10 min and then quenched with 0.125 M glycine for 5 min at room temperature. After returning to normal conditions (21% O2), cells were washed with PBS and incubated in 3% BSA (Sigma) with CD16/CD32 Fc Block™ (1:250, BD Pharmingen, Cat#553142) for 15 min. Next, the anti-CD45-PE antibody (Clone 30-F11, 1:400, BD Pharmingen, Cat#553081) was added for 40 min, followed by washes with PBS. CD45+ or GFP+ cells were sorted into 1% BSA in PBS using Cell Sorter BD FACSAriaII. Subsequently, 50,000 cells were lysed in cold lysis buffer (10 mM Tris-HCl, pH 7.4, 10 mM NaCl, 3 mM MgCl2 and 0.1% IGEPAL CA- 630) for 5 min on ice. Cells were then centrifuged at 500xg for 8min, and cell pellets were resuspended in transposition reaction according to the standard ATACseq protocol using 2.5 ul of Tn5 enzyme per 50 000 of cells ^73^. Cells were then centrifuged and reverse crosslinked (50 mM Tris-Cl, 1 mM EDTA, 1% SDS, 0.2 M NaCl, 200 μg/ml proteinase K) at 65°C with an overnight shaking at 1200 rpm. Zymo DNA Clean & Concentrator-5 columns (ZymoResearch) were used to purify DNA and the sequencing libraries were prepared according to the original ATACseq protocol ^73^. The ATACseq libraries were assessed for appropriate quality using a Bioanalyzer 2100 and subjected to paired-end sequencing (2x151bp) using NovaSeq 6000 (Illumina) at the Laboratory of Sequencing (Nencki Institute). Two independent biological replicates were analysed per each condition. The ATACseq data generated in this project is available at the NCBI platform under GSE279538 accession number.

### ATACseq data processing and analysis

The quality of raw fastq data was assessed using FASTQC software (https://www.bioinformatics.babraham.ac.uk/projects/fastqc/). Reads were trimmed using Trimmomatic (version 0.36) to remove adapter, transposase sequence short reads, and low quality 5′ and 3′ bases ^65^. Paired-end reads were then aligned to mm10 reference genome using Hisat2 (version 2.1.0) with parameters -X 2000 ^74^. Duplicate reads were subsequently removed with Picard’s MarkDuplicates tool (version 2.26.2, https://broadinstitute.github.io/picard/), and reads mapping to the mitochondrial genome were also excluded. Only properly paired and uniquely mapped reads were kept for downstream analysis using samtools view with the options -q 30 -f 2 -F 256. Peaks were called using MACS2 (version 2.1.1.2) with parameters set to -f BAPME -q 0.01 –nomodel –shift 0 ^75^. The obtained peaks were then filtered for peaks overlapping mm10 ENCODE blacklisted genomic regions. To visualize the overlap of open genomic regions (peaks) from different conditions, the makeVennDiagram function from the ChIPpeakAnno package was used with minoverlap parameter set to 200 ^76^. All sequencing tracks were generated by IGV using a normalized bigWig file created with deepTools (bamCoverage) and normalized to genome coverage – RPGC ^71,72^. Changes in chromatin accessibility were assessed using edgeR with model that includes all groups ^68^. The following steps were performed to create a set of consensus ATAC peaks for the read count matrix. First, peaks for two replicates in each condition were intersected and only peaks with overlap > 200 bp were included. Then obtained peaks from all conditions were merged with reduction of overlapping regions using the “reduce” function from the GenomicRanges package to generate consensus peak lists ^77^. Peaks with FDR < 0.05 (Benjamini and Hochberg - corrected P-value) and |log2 fold change| >= 0.6 were classified as significantly different.

### Annotation for differential ATACseq peaks to genomic features

The peaks were annotated to genomic features using ChIPseeker, with the promoter region defined as +/- 2,000 bp around the transcription start site (TSS) and the overlap parameter set to ’all’ to annotate peaks to the nearest gene ^78^. Each peak was annotated to only one genomic feature according to the default annotation priority in ChIPseeker. To assign promoter peaks to all possible genes, a custom-written script in R was used based on biomaRt ^77,79^ and GenomicRanges packages ^79^. Functional enrichment analysis of differentially accessible regions nearby genes was performed using enrichGO function from clusterProfiler ^69^. Revigo tool was used to merge similar GO terms and visualisation ^70^. Motif enrichment analysis for selected ATAC-seq peaks was performed using HOMER findMotifsGenome.pl (http://homer.ucsd.edu/homer/motif/) for known TF motifs from HOMER custom database with the option -size given. For local motif analysis in ATAC-seq peaks, FIMO from the MEME suite was used ^80^. For the figures, enriched motifs were shown after further filtering to include only those motifs with q-value < 0.01 and where the expression of the transcription factor gene was sufficiently high, defined as a sum of the log2- transformed CPM values across the samples of at least 2.

## AVAILABILITY OF DATA AND MATERIALS

All sequencing data generated in this project are available at the NCBI platform (https://www.ncbi.nlm.nih.gov/) under GSE279536 (RNAseq) and GSE279538 (ATACseq) accession numbers.

## COMPETING INTERESTS

The authors declare no competing interests.

## AUTHORS CONTRIBUTIONS

MD co-designed and performed the experiments, interpreted the results and co-wrote the manuscript. PR, BK (Kaza), SL, SC, AK, TO and KP performed the experiments. AEM, ARM and JM provided experimental guidance on the project and co-wrote the manuscript. BK (Kaminska) provided guidance on the project, infrastructure for the experiments and co-wrote the manuscript. KBL conceived the project and secured the funding; designed, performed, and supervised the experiments; interpreted the results, and wrote the manuscript. All authors commented on and approved the manuscript.

## ACKNOWLEDGMENTS

We thank Pawel Segit, Karol Jacek and Adria-Jaume Roura Canalda for their insightful suggestions regarding the bioinformatic analyses. We thank Bartek Gielniewski and Paulina Szadkowska from the Sequencing Core Facility at the Nencki Institute for sequencing the samples.

## FUNDING

This work was funded by the National Science Center grant (Poland) 2019/33/B/NZ1/01556 (awarded to KBL).

## Notes

### Competing Interest Statement

The authors have declared no competing interest.

### Summary of Updates

We have included some new data validating the hypoxia-induced properties of GAMs in promoting glioma invasion (in Fig. 4). This resulted in addition of a new author, who performed this work. We have updated the discussion and improved the manuscript text to clarify our results. Labels on some graphs have been improved for a better visibility. We have improved some schemes in figures/supplementary figures and adjusted scales in the scRNAseq UMAP plots for a better clarity of the results (Fig 1 and Suppl. Fig 2).

