## Supplementary text for "Hypoxic stress dysregulates functions of glioma-associated myeloid cells through epigenomic and transcriptional programs"

### Table of contents

Supplementary Figure legends Page 2

Supplementary Table legends Page 7

Primer sequences Page 8

### SUPPLMENTARY FIGURE LEGENDS

**Supplementary Figure 1. A mild hypoxia and a hypoxia-mimetic drug dysregulates the expression of *Lgals3, P2ry12* and *Tmem119* in BV2 cells.**

**A,** Expression of *Lgals3* in BV2 cells exposed to 1% $O_2$ or $CoCl_2$ for 16 hours was analysed with qPCR in relation to *Rn18s* housekeeping gene. *Vegfa* was used as hypoxia control. The mean fold change with standard deviation (SD) is shown from three independent biological replicates. Two-tailed student t-test determined the statistical significance (* $p$<0.05, **** $p$<0.0001).

**B,** BV2 and RAW 264.7 cell lines were treated as in (**A**) and subjected to western blotting with anti-LGALS3 antibody. HIF-1α was used as a positive control for hypoxia. A representative experiment of three independent repeats is shown.

**C,** Expression analysis of *P2ry12 and Tmem119* from cells in (**A**), as indicated, in relation to *Rn18s* housekeeping gene. The mean fold change with standard deviation (SD) is shown from three independent biological replicates.

**Supplementary Figure 2. Hypoxia dysregulates the expression of *LGALS3, P2RY12* and *TMEM119* in human GBM samples *in vivo*.**

**A,** UMAP plot showing clusters of monocytes and TAM subpopulations from human newly diagnosed GBM tumours as identified in the scRNA-seq Antunes-dataset (GSE163120) [6].

**B,** UMAP plot showing the expression of "hypoxia score" (see Supplementary Table 1) in the dataset from (**A**) [6].

**C-E,** UMAP plots showing the expression of (**C**) *LGALS3*, (**D**) *TMEM119* and (**E**) *P2RY12* in the dataset from (**A**) [6].

**F,** UMAP plot showing clusters of monocytes and TAM subpopulations from human recurrent GBM tumours, as identified in the scRNA-seq Antunes-dataset (GSE163120) [6].

**G,** UMAP plot showing the expression of "hypoxia score" in the dataset from (**F**) [6].

**H-J,** UMAP plots showing the expression of (**H**) *LGALS3*, (**I**) *TMEM119* and (**J**) *P2RY12* in the dataset from (**F**).

**Supplementary Figure 3. Quality check of RNA samples from BV2-GL261 co-cultures for RNAseq after glyoxal fixation and cell sorting**

**A,** Schematic representation of cell processing from glioma-microglia co-culture samples for flow cytometry and subsequent RNA sequencing.

**B,** Gating strategy to sort BV2 or GL261 cells from co-culture experiment, where microglia cells were defined as $CD45^+$ and glioma cells as $GFP^+$.

**C,** Bioanalyzer profiles of total RNA derived from BV2 cells after glyoxal fixation. 5 ng of total RNA was separated on RNA pico chips using Agilent 2100 Bioanalyzer.

**D,** Expression analysis (qPCR) of *Vegfa* and *Glut1* in BV2 cells after glyoxal fixation, as indicated in relation to *Rn18s* housekeeping gene.

**Supplementary Figure 4. Genome-wide transcriptomic analysis in BV2 microglial cells treated with glioma co-culture and hypoxia.**

**A-C,** Volcano plots showing expression changes between glioma co-cultured (CC) microglial cells and microglia monocultures (Ctrl) **(A)**, glioma co-cultured (CC) microglial cells in hypoxia (H) and normoxia (N) **(B)**, microglia monocultures exposed to hypoxia or normoxia

**(C)**. Each point on a plot corresponds to one gene. Genes with significantly altered expression levels (adj. P-value<0.05 and |log fold change |≥1) were marked in blue. The numbers of genes significantly down- or upregulated are written above the plots.

**D-F**, Average gene expression levels in the sets of up- or downregulated genes in glioma co-cultured microglial cells (CC) compared to microglial monocultures (Ctrl) **(D)**, glioma co-cultured microglial cells (CC) in hypoxia compared to normoxia **(E)** and in microglia monocultures (Ctrl) exposed to hypoxia or normoxia **(F)**.

**G,** HIF metagene expression in BV2 cells treated with hypoxia as monocultures (Ctrl) or co-cultures with glioma (CC). HIF metagene set was defined by Lombardi et al., [26]. The average gene expression signal of the HIF metagene score is represented as read counts per million. Two-way ANOVA and post-hoc Tukey test determined the statistical significance.

**H,** Transcription factor (TF) motif enrichment analysis using HOMER tool on the promoters of significantly upregulated genes in BV2 cells after hypoxic co-culture with glioma cells in relation to normoxic co-culture.

**Supplementary Figure 5. Epigenetic analysis of BV2 microglial cells in response to glioma co-culture and hypoxic treatment.**

**A,** Immunofluorescent staining of H3K27me3 (red) and nuclei (DAPI, blue) was performed in BV2-GL261 co-cultures in hypoxia (Hyp) and normoxia (Norm). Representative images show BV2 (microglia) and GL261 (glioma GFP+, green) cells. Scale bar, 20 μm.

**B,** Quantitation of nuclear fluorescence intensity of H3K27me3 staining from (A) was preformed and five different fields of view were analysed in each condition. Unpaired two-tailed student's t test was used to determine statistical significance (**** $P < 0.0001$). A representative experiment of three independent biological repeats is shown.

**C,** Immunofluorescent staining of H3K9me3 (red) and nuclei (DAPI, blue) was performed in BV2-GL261 co-cultures in Hyp and Norm. Representative images show BV2 and GFP-GL261 cells. Scale bar, 20 µm.

**D,** Quantitation of nuclear fluorescence intensity of H3K9me3 staining from (**C**) was preformed and five different fields of view were analysed in each condition. Unpaired Student's t test was used to determine statistical significance (**** $P < 0.0001$). A representative experiment of three independent biological repeats is shown.

**E,** Schematic representation of cell processing from glioma-microglia co-culture samples for flow cytometry and subsequent ATACseq experiment.

**F**, Venn diagram showing specific or common ATACseq peaks detected in all conditions tested, as indicated. The percentage of peaks in the total amount of peaks is shown in brackets for each condition.

**G-I**, Volcano plots showing differentially altered ATACseq peaks in glioma co-cultured (CC) microglial cells versus microglia monocultures **(G)**, glioma co-cultured microglial cells (CC) in hypoxia versus normoxia **(H)**, microglia monocultures in hypoxia versus normoxia **(I)**. Each point on the plot corresponds to one peak. Statistically significant peaks with FDR < 0.05 and |log2 fold change| ≥ 0.6 are marked in blue. The numbers of chromatin accessibility peaks significantly increased or decreased are written above the plots.

**Supplementary Figure 6. Changes in chromatin accessibility enhanced by combined glioma culture in hypoxic treatment when compared to hypoxia alone.**

**A,** A heatmap showing z-scores for peaks increased (red) or decreased (dark blue) in BV2 cells co-cultured (CC) with glioma cells. Peaks which were additionally increased or decreased in CC versus Ctrl hypoxic BV2 cells are projected in magenta or pale blue, respectively.

**B,** A pie chart showing differentially regulated ATACseq peaks from (**A**) annotated to specific genomic regions, as indicated in the figure legend.

**C,** Overrepresentation analysis showing GO processes in peaks which were additionally decreased or increased in CC versus Ctrl hypoxic BV2 promoter regions. The top 15 GO processes were selected based on the lowest adjusted p-values; closely related terms were merged using REVIGO. The colour gradient reflects the gene count in each associated process.

**D,** A volcano plot showing differentially altered ATACseq peaks occurring at promoter regions in glioma co-cultured (CC) microglial cells in hypoxia versus normoxia (numbers written above the plot). Each point on the plot corresponds to one peak. Statistically significant peaks are defined as FDR $< 0.05$ and $|\log2 \text{ fold change}| \geq 0.6$. Additionally, the promoter peaks that were even more decreased or increased in CC versus Ctrl in hypoxic conditions are projected in pale blue or magenta, respectively. Examples of genes associated with those peaks are written below the plot.

**E,** Overrepresentation analysis showing GO processes in ATACseq peaks which were additionally decreased or increased in CC versus Ctrl hypoxic conditions and occurred at intron and exon regions. The top 15 GO processes were selected based on the lowest adjusted p-values, and closely related terms were merged using REVIGO. The colour gradient reflects the gene count in each associated process.

**Supplementary Figure 7. Concordant changes in gene expression and chromatin accessibility in response to hypoxia in BV2 cells co-cultured with glioma.**

**A-C,** Overrepresentation analysis showing GO processes in concordant (mRNA and chromatin accessibility changes) upregulated (**A**) or downregulated (**B**) genes in BV2 cells co-cultured with glioma cells in hypoxic conditions. In (**C**) are shown processes that were downregulated in hypoxic co-cultures (CC Hyp) and additionally downregulated in hypoxic co-cultures in comparison to hypoxic monocultures (CC Hyp < Ctrl Hyp). The box shows the genes that were most often occurring in identified GO processes in (**C**).

### SUPPLEMENTARY TABLE LEGENDS

**Supplementary Table 1. "Hypoxia score" genes, related to Figure 1.**

Top genes were extracted from publicly available scRNAseq datasets [8,6].

**Supplementary Table 2. RNAseq GO pathways analysis in BV2 cells in CC under hypoxia.**

Overrepresentation analysis (ORA) of GO processes enriched in BV2 at the differentially regulated genes in hypoxic co-cultures with glioma. In separate tabs are shown GO processes or differentially expressed genes, as indicated.

**Supplementary Table 3. TAM subcluster marker genes, related to Figure 3.**

Top marker genes defined for TAM subclusters were extracted from publicly available scRNAseq datasets [5-8].

**Supplementary Table 4. RNAseq GO pathways analysis in GL261 cells in CC or Ctrl under hypoxia.**

Overrepresentation analysis of GO processes enriched in GL261 cells at the differentially upregulated genes in hypoxic co-cultures with microglia cells or in hypoxic monocultures of GL261. In separate tabs are shown GO processes or differentially expressed genes, as indicated.

**Supplementary Table 5. ATACseq GO pathway analysis in BV2 cells in CC or Ctrl under hypoxia.**

Overrepresentation analysis of GO processes for genes with differentially altered ATACseq peaks in genic region in BV2 cells co-cultured with glioma cells under hypoxia. In separate tabs are shown GO processes related to identified peaks or differentially regulated ATACseq peaks assigned to genes, as indicated.

**Supplementary Table 6. GO pathway analysis in BV2 cells in CC or Ctrl under hypoxia.**

Overrepresentation analysis of GO processes enriched in BV2 cells at the differentially regulated genes with concordant changes in chromatin accessibility in hypoxic co-cultures with glioma cells.

**Supplementary Table 7. FIMO analysis for peaks in myeloid marker genes related to Figure 6D-G.**

TF motifs within the changing ATACseq peaks in *Gpnmb, Lgals3, P2ry12 and Olfml3* genes were identified using FIMO tool.

**Supplementary Table 8. Correlation data related to Figure 6H-J.**

Correlations between expression of the "hypoxia score" and selected hypoxia-regulated TAM marker genes in scRNAseq datasets for TAMs in mouse and human GBM.

### PRIMER SEQUENCES USED IN QPCR ANALYSIS

*Rn18s* F: CGGACATCTAAGGGCATCACA
*Rn18s* R: AACGAACGAGACTCTGGCATG
*Lgals3* F: AACACGAAGCAGGACAATAACTGG
*Lgals3* R: GCAGTAGGTGAGCATCGTTGAC
*Vegfa* F: GTCCGATTGAGACCCTGGTG
*Vegfa* R: GCTGGCTTTGGTGAGGTTTG
*Glut1* F: ATCCCATCCACCACACTCAC
*Glut1* R: GAGAAGCCCATAAGCACAGC
*P2ry12* F: CAAGGGGTGGCATCTACCTG
*P2ry12* R: GCCTTGAGTGTTTCTGTAGGGT
*Tmem119* F: ACTACCCATCCTCGTTCCCTGA
*Tmem119* R: TAGCAGCCAGAATGTCAGCCTG
