## Supplementary Figures for "Hypoxic stress dysregulates functions of glioma-associated myeloid cells through epigenomic and transcriptional programs"

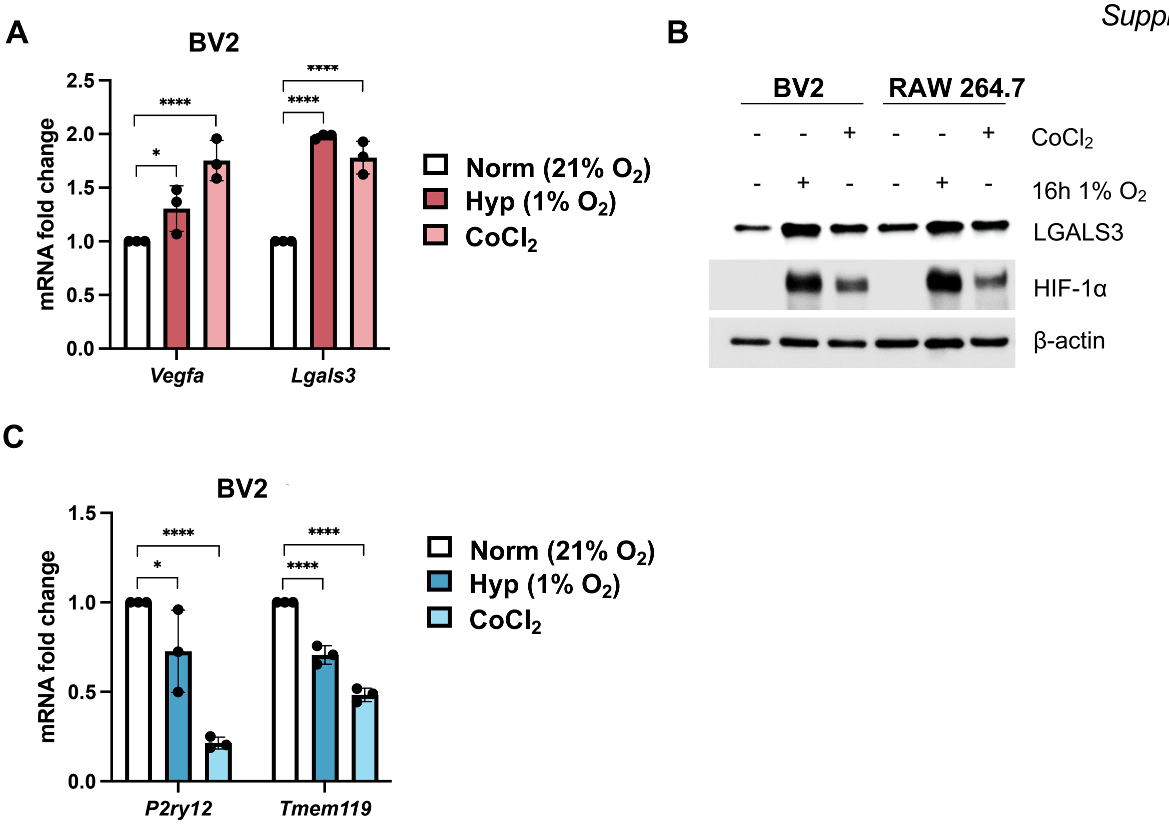
A
BV2
mRNA fold change
2.5
2.0
1.5
1.0
0.5
0.0
*
****
****
****
Vegfa
Lgals3
Norm (21% O2)
Hyp (1% O2)
CoCl2
B
BV2
RAW 264.7
CoCl2
16h 1% O2
LGALS3
HIF-1α
β-actin
C
BV2
mRNA fold change
1.5
1.0
0.5
0.0
*
****
****
****
P2ry12
Tmem119
Norm (21% O2)
Hyp (1% O2)
CoCl2

*Supplementary Figure 2*

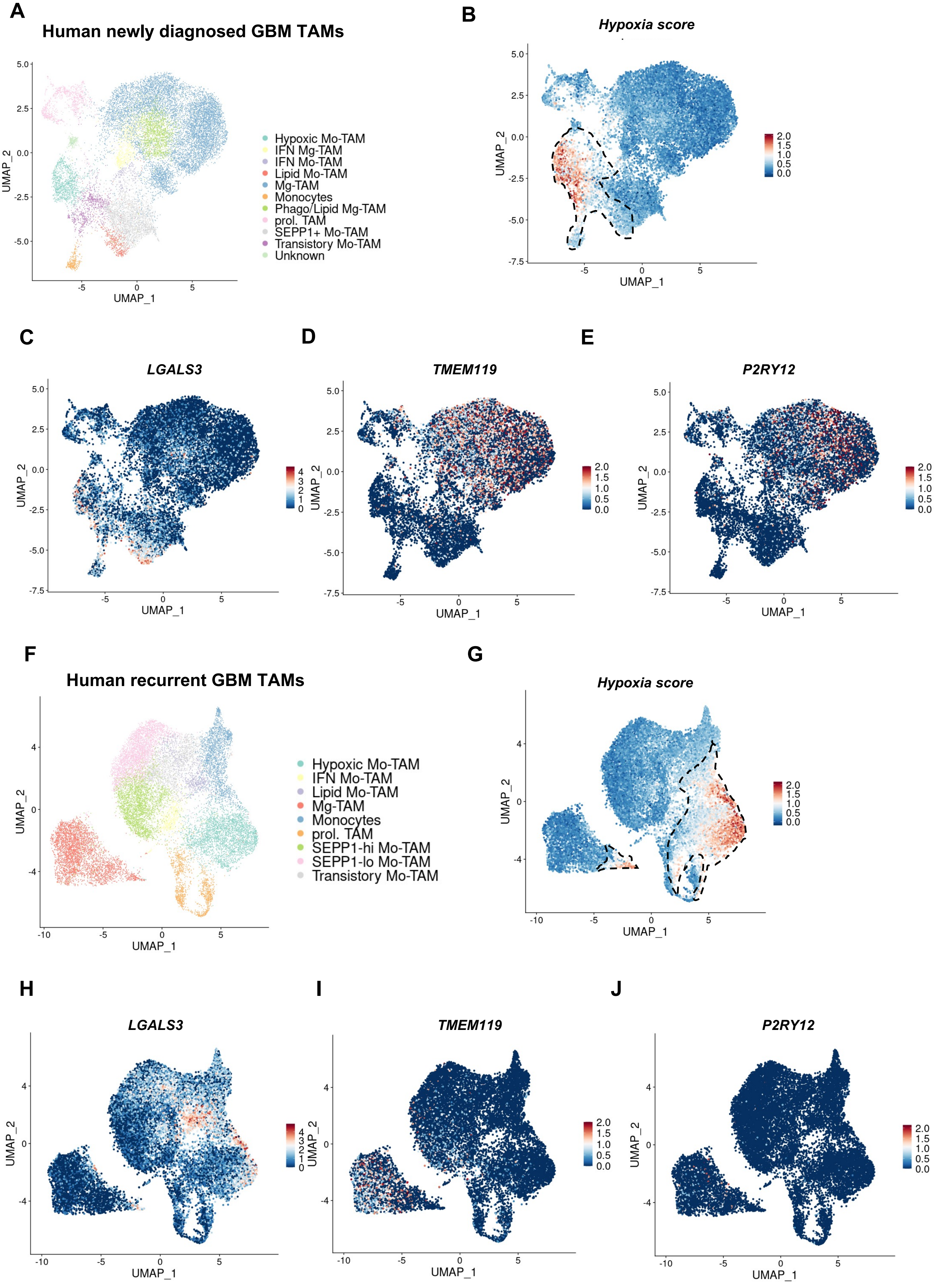

*Supplementary Figure 3*

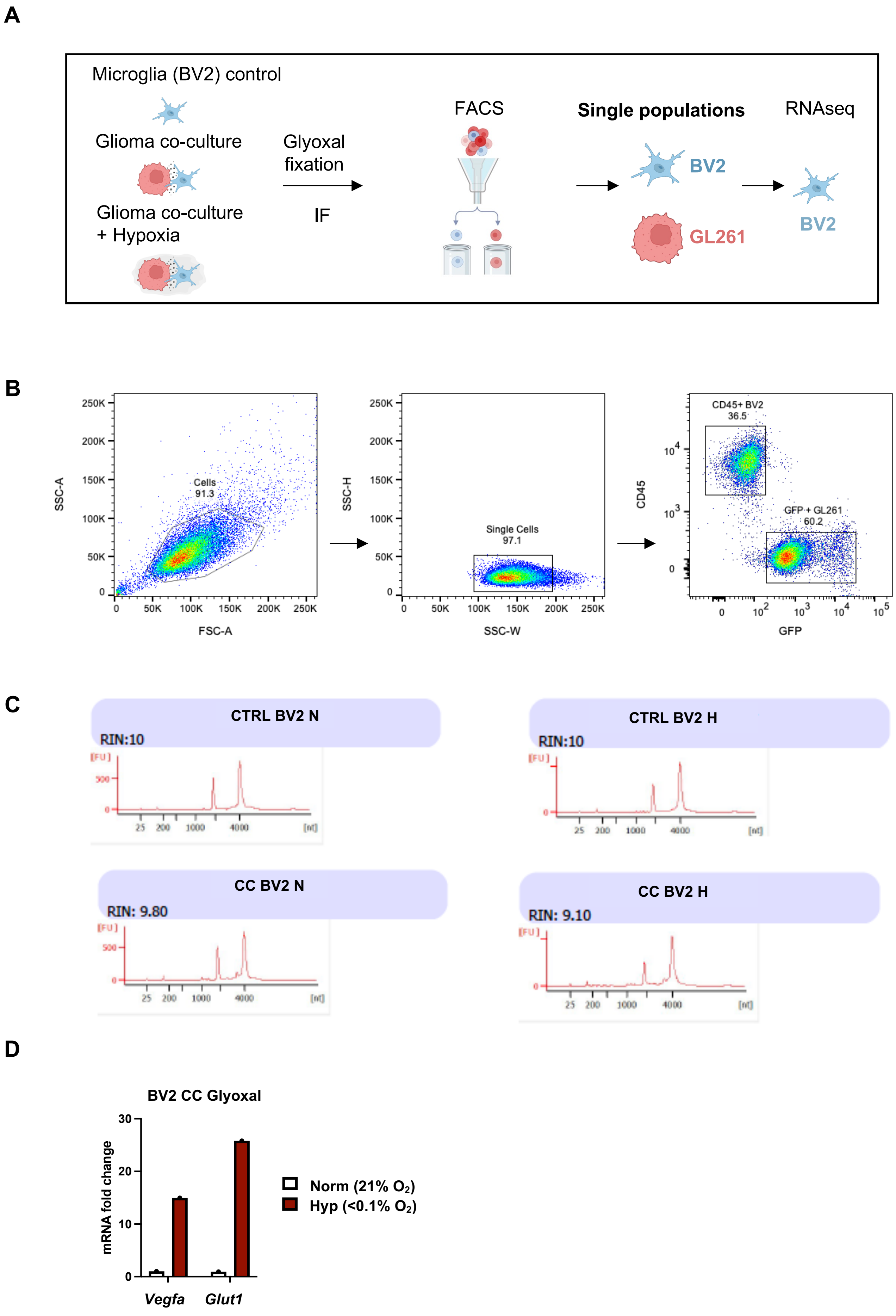

*Supplementary Figure 4*

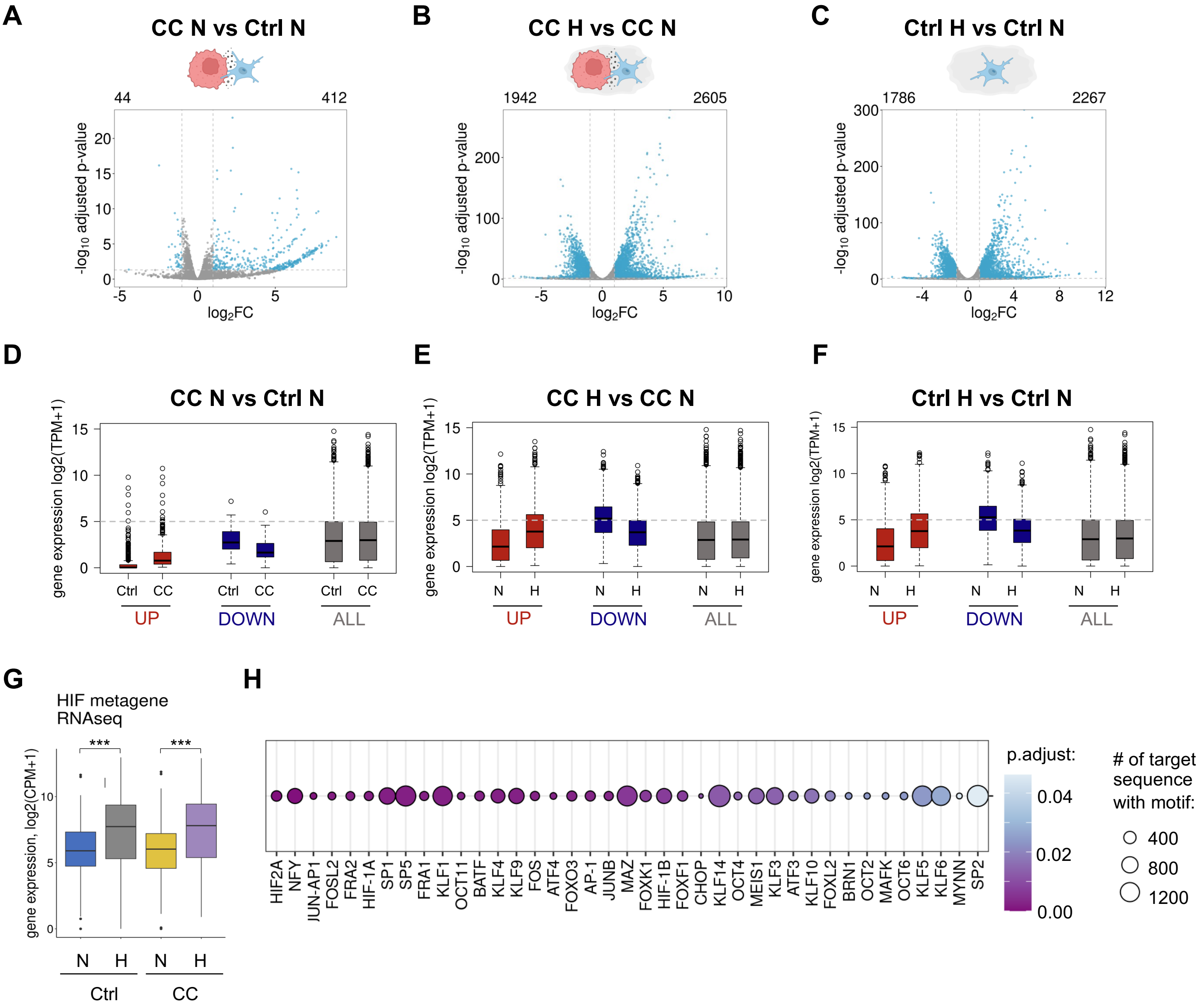

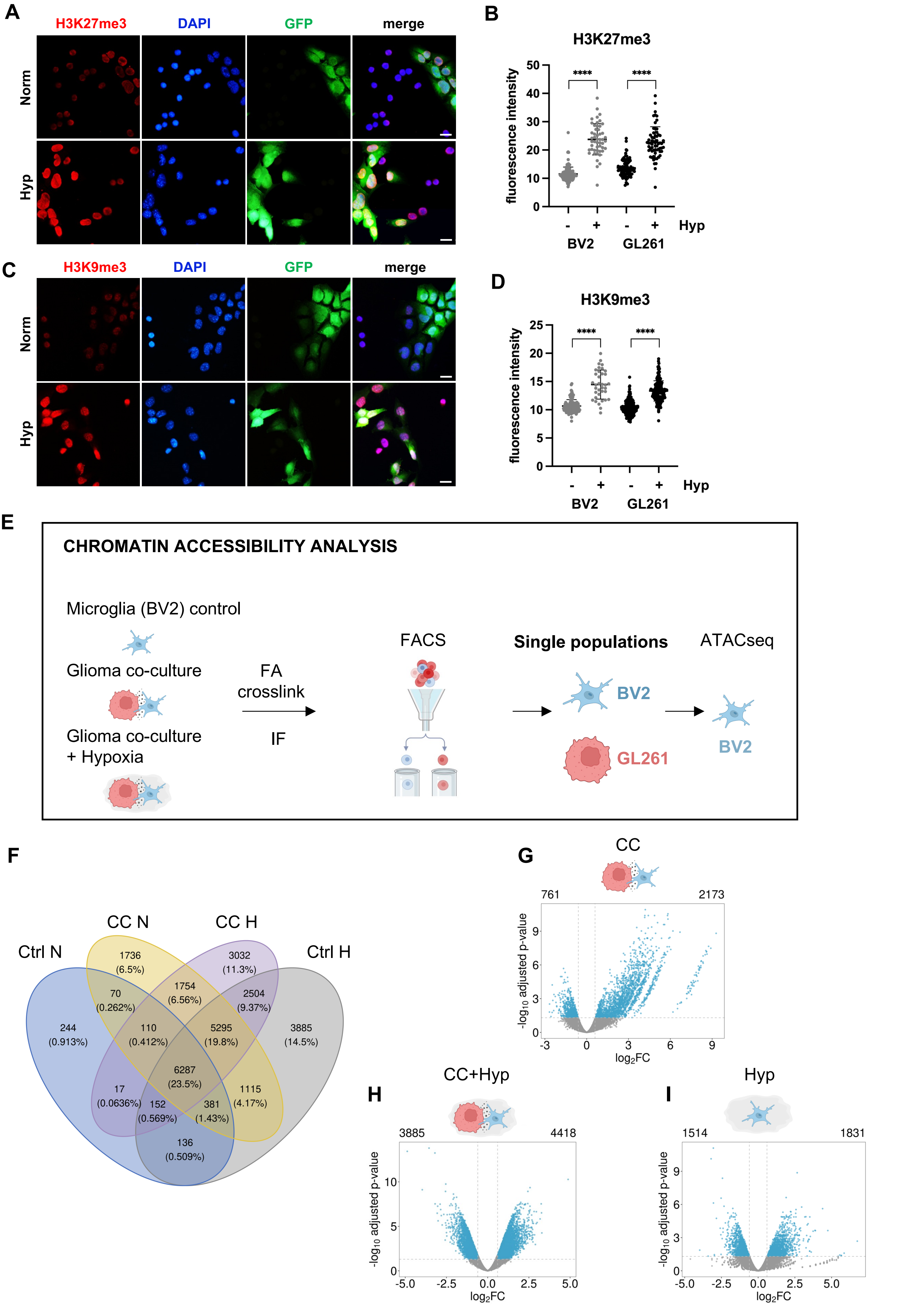
A
H3K27me3
DAPI
GFP
merge
Norm
Hyp
B
H3K27me3
fluorescence intensity
****
****
- + - + Hyp
BV2 GL261
C
H3K9me3
DAPI
GFP
merge
Norm
Hyp
D
H3K9me3
fluorescence intensity
****
****
- + - + Hyp
BV2 GL261
E
CHROMATIN ACCESSIBILITY ANALYSIS
Microglia (BV2) control
Glioma co-culture
Glioma co-culture + Hypoxia
FA crosslink
IF
FACS
Single populations
BV2
GL261
ATACseq
BV2
F
Ctrl N
CC N
CC H
Ctrl H
244 (0.913%)
1736 (6.5%)
3032 (11.3%)
3885 (14.5%)
70 (0.262%)
1754 (6.56%)
2504 (9.37%)
110 (0.412%)
5295 (19.8%)
6287 (23.5%)
17 (0.0636%)
152 (0.569%)
381 (1.43%)
1115 (4.17%)
136 (0.509%)
G
CC
761
2173
-log10 adjusted p-value
log2FC
H
CC+Hyp
3885
4418
-log10 adjusted p-value
log2FC
I
Hyp
1514
1831
-log10 adjusted p-value
log2FC

*Supplementary Figure 6*

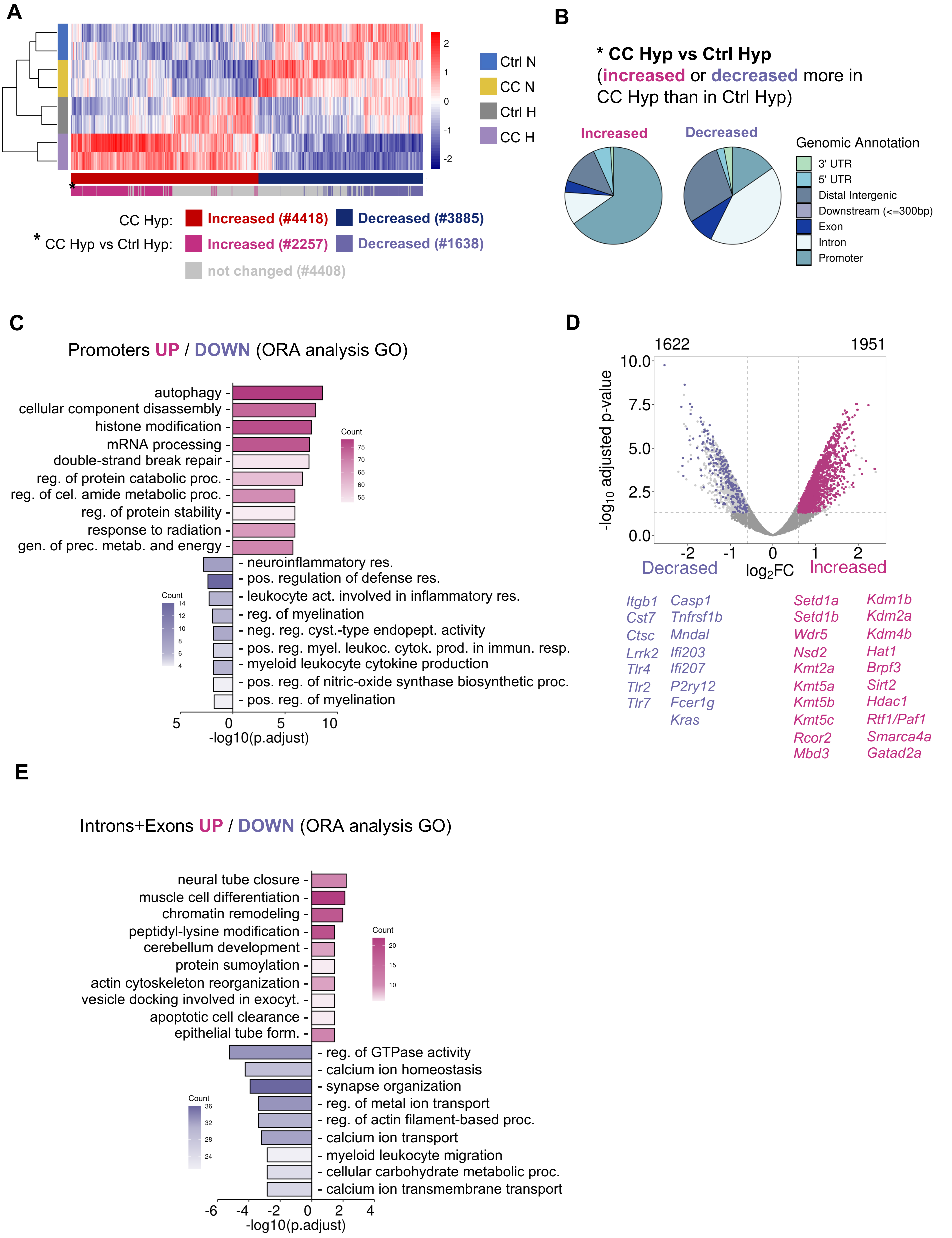

*Supplementary Figure 7*

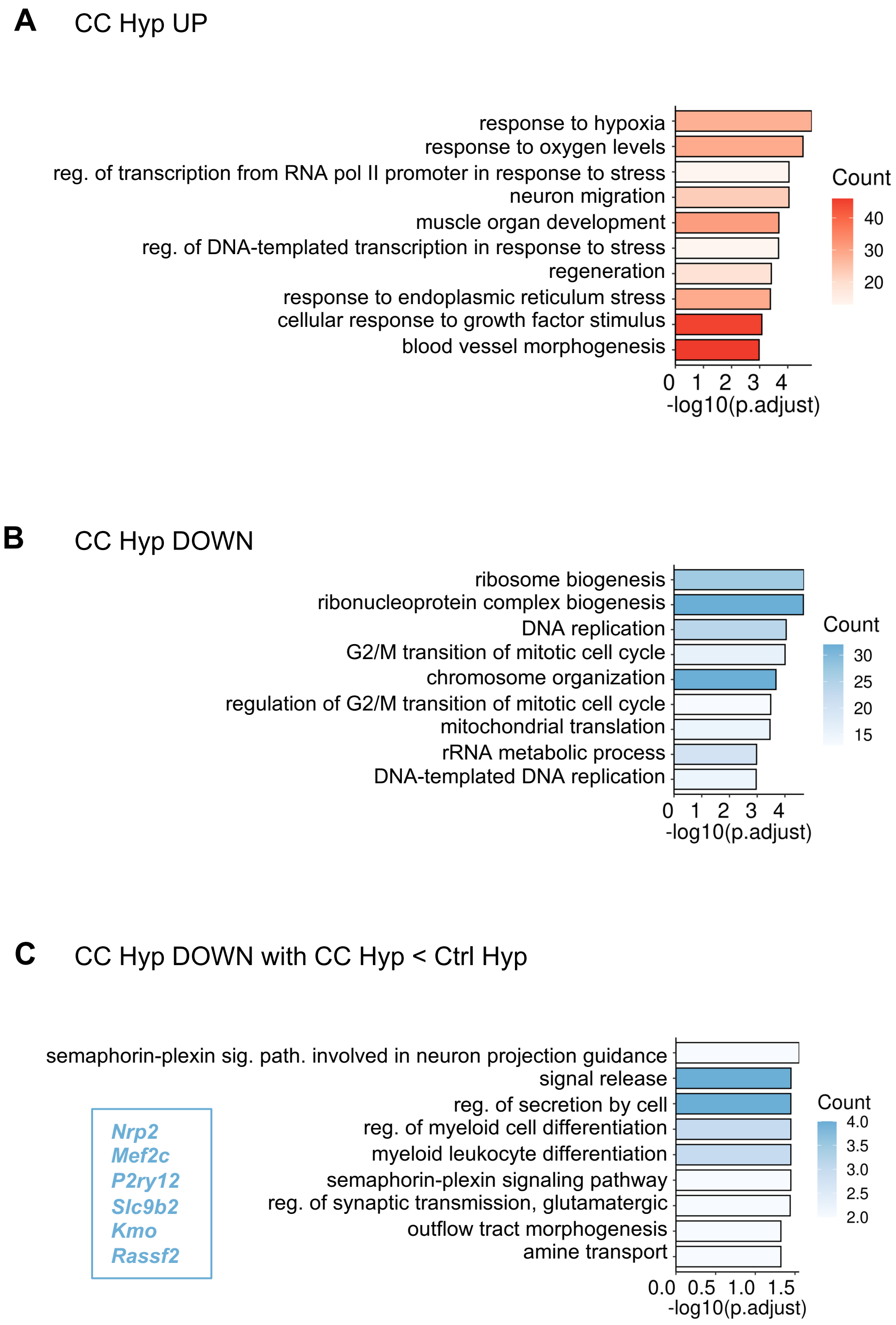
